# The Ebola virus VP40 matrix undergoes endosomal disassembly essential for membrane fusion

**DOI:** 10.1101/2022.08.24.505067

**Authors:** Sophie L. Winter, Gonen Golani, Fabio Lolicato, Melina Vallbracht, Keerthihan Thiyagarajah, Samy Sid Ahmed, Christian Lüchtenborg, Oliver T. Fackler, Britta Brügger, Thomas Hoenen, Walter Nickel, Ulrich S. Schwarz, Petr Chlanda

## Abstract

Ebola viruses (EBOVs) are filamentous particles, whose shape and stability are determined by the VP40 matrix. Virus entry into host cells occurs via membrane fusion in late endosomes; however, the mechanism of how the remarkably long virions undergo uncoating including virion disassembly and nucleocapsid release into the cytosol, remains unknown. Here, we investigate the structural architecture of EBOVs entering host cells and discover that the VP40 matrix disassembles prior to membrane fusion. We reveal that VP40 disassembly is caused by the weakening of VP40-lipid interactions driven by low endosomal pH that equilibrates passively across the viral envelope without a dedicated ion channel. We further show that viral membrane fusion depends on VP40 matrix integrity, and its disassembly reduces the energy barrier for fusion stalk formation. Thus, pH-driven structural remodeling of the VP40 matrix acts as a molecular switch coupling viral matrix uncoating to membrane fusion during EBOV entry.

## Introduction

Ebola viruses (EBOVs) are highly pathogenic negative-sense RNA viruses causing severe outbreaks of viral hemorrhagic fever in humans with high case fatality rates^1^. They enter host cells by macropinocytosis and undergo cytosolic entry in late endosomal compartments, where the fusion of the viral and endosomal membranes leads to genome release into the cytoplasm. EBOVs are characterized by their filamentous morphology which is determined by the viral matrix protein VP40 that drives budding of virions reaching up to several micrometers in length^2,3^. VP40 interacts with negatively charged lipids^4–6^ to assemble into a quasi-helical scaffold underneath the viral membrane^7,8^ and is critical for the incorporation of the viral nucleocapsid into the virions by so far unknown VP40-nucleocapsid interactions. The EBOV nucleocapsid is composed of the nucleoprotein (NP), viral protein (VP)24, and VP35^3,9,10^ which together encapsidate the single-stranded RNA genome. Upon host cell entry, the nucleocapsid needs to dissociate from the virus particle and viral genome to enter the cytoplasm and enable genome replication and transcription^11^. These processes together are referred to as virus uncoating, which involves the weakening of protein-protein and protein-membrane interactions inside the virus lumen. The resulting changes in virion architecture allow the timely nucleocapsid release upon membrane fusion^12^. It is well established that fusion of the viral and endosomal membrane relies on interactions with the EBOV fusion protein GP, which is the only transmembrane protein that studs the viral envelope^13–15^. GP-mediated membrane fusion is triggered after proteolytic processing of GP by host cell cathepsin proteases^16^ and depends on the interaction of the cleaved GP subunit GP1 with the late endosomal Niemann-Pick C1 (NPC1) receptor^17–20^. However, the molecular mechanism of how the remarkably long EBOVs undergo uncoating during cytosolic entry remains enigmatic. A growing body of evidence shows that matrix disassembly during viral entry can trigger a cascade of events required for viral uncoating and efficient virus entry^21,22^. While the structure of isolated Ebola virions is well characterized, it is currently unknown whether the VP40 matrix undergoes conformational changes during virion entry and factors initiating EBOV disassembly remain to be elucidated. In addition, a mechanistic understanding of how interactions between the EBOV VP40 matrix, the viral membrane and nucleocapsid are modulated during viral entry is still missing. Since EBOVs belong to the late-penetrating viruses, which require low endosomal pH for cytosolic entry^23^, the acidic environment may serve as one of the triggers for virion uncoating.

Here, we investigate EBOV uncoating and the role of VP40 during virus entry into host cells by characterizing EBOVs in endosome-mimicking conditions *in vitro* and in endo-lysosomal compartments by *in situ* cryo-electron tomography (cryo-ET), which is complemented by membrane modelling approaches, lipidomics, and time-lapse fluorescence imaging. We find that the VP40 matrix and its interactions with lipids in the viral envelope are sensitive to low pH, which passively equilibrates across the viral envelope in acidic environments. This leads to the disassembly of the matrix layer allowing for fusion and genome release.

## Results

### The Ebola virus VP40 matrix undergoes disassembly in endosomal compartments

To shed light on EBOV endosomal uncoating at molecular resolution, we infected Huh7 cells cultured on electron microscopy grids with EBOVs (strain Mayinga) in BSL4 containment. Infected cells were chemically fixed after multiple rounds of infection had occurred at 22 or 48 h post-infection (Fig. 1 A, Fig. S1–3). After vitrification and cryo-focused ion beam (FIB) milling of the infected cells, we performed *in situ* cryo-ET of endosomal compartments containing EBOV particles (Fig. 1 B-G, Supplementary Video 1). Late endosomal compartments were identified by the presence of vesicles and membrane fragments (white arrow, Fig. 1 F), which are likely products of lysosomal degradation. In addition, we observed the accumulation of crystalline lipidic structures with a spacing of 3.2 nm (Fig. S1), consistent with the spacing found in cholesterol ester crystals previously described in lamellar bodies, lipid droplets, and isolated low-density lipoprotein particles^24–26^. Interestingly, Ebola virions in late endosomes retained their filamentous morphology and displayed well-defined nucleocapsids of approximately 20 nm in diameter (Fig. 1, Fig. S 2). They appeared condensed and resembled nucleocapsid structures formed by truncated EBOV NP alone^27^ but lacked the regular protrusions observed in nucleocapsids of isolated virions (Fig. S3). However, the VP40 matrix layer was detached from the envelope as apparent from the empty space adjacent to the EBOV membrane and disordered protein densities, which presumably represent disassembled VP40, were surrounding the nucleocapsid in the EBOV lumen (Fig. 1 D-G). Importantly, none of the five EBOV captured in endosomes displayed ordered VP40 matrices, and two virions had engulfed intraluminal vesicles (Fig. S2). In contrast, budding virions and extracellular virions adjacent to the plasma membrane of infected cells displayed assembled VP40 layers with VP40 proteins visible as distinct densities lining the membrane (Fig. 5 H-K, S4, n=8), similar to the VP40 layer in isolated virions (Fig. S3). Overall, this data indicates that EBOV uncoating involves VP40 disassembly in late endosomal compartments and suggests that endosomal VP40 disassembly occurs prior to GP-mediated membrane fusion.

**Figure 1:**
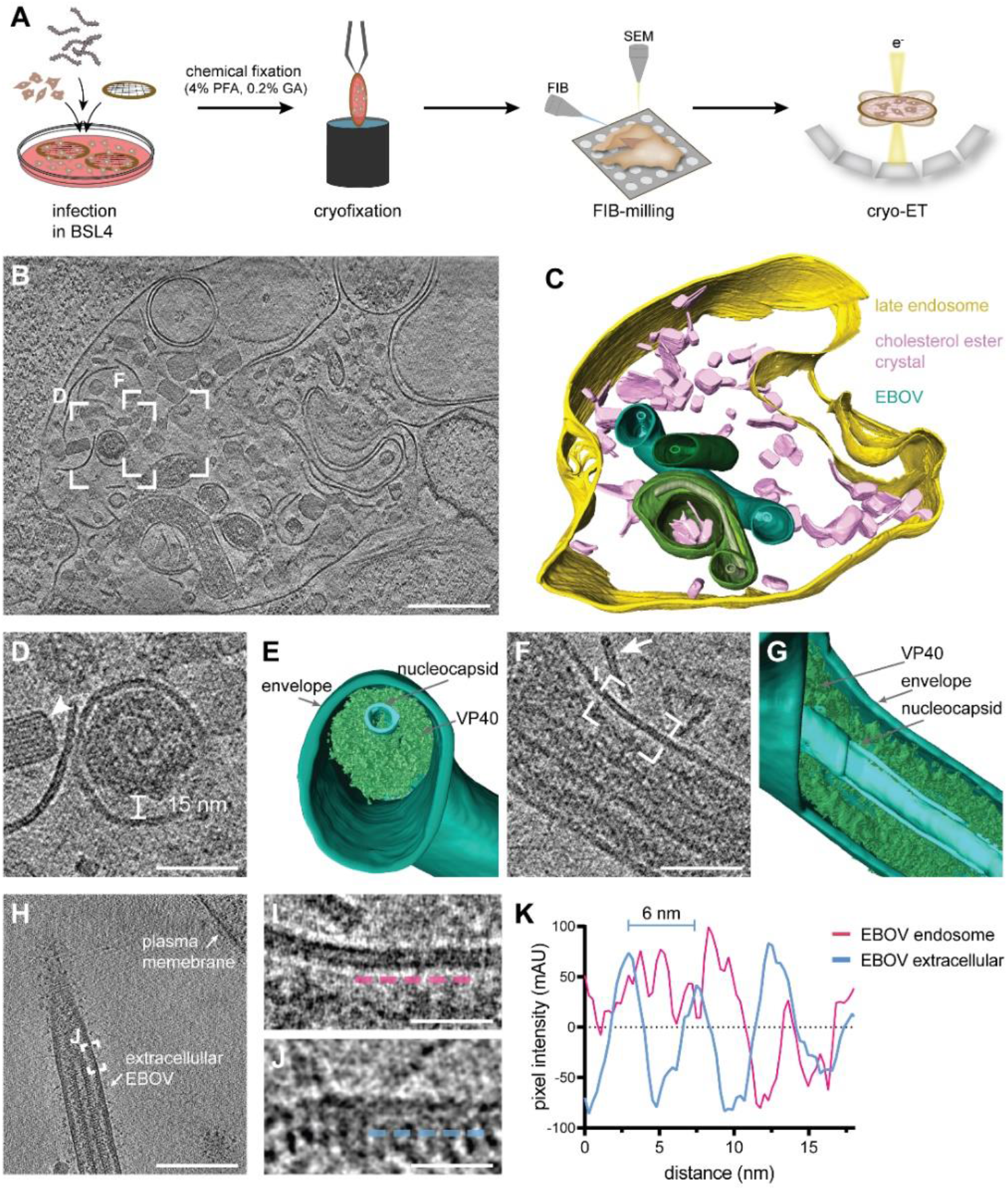
*In situ* cryo-electron tomography of Ebola virions localized in endosomes of an infected cell. **(A)** Schematic of the *in situ* cryo-ET workflow, including infection of cells grown on electron microscopy grids and chemical fixation for biosafety reasons before removal from BSL4. Cryofixation was performed prior to cell thinning by cryo-FIB milling and imaging by cryo-ET. **(B)** Slice through a tomogram showing EBOVs inside a late endosomal compartment. **(C)** 3D segmentation of the delimiting membrane (yellow), cholesterol ester crystals (pink), viral membranes (green) and nucleocapsids (light green) for visualization. **(D)** Magnified view of the area highlighted in (B) showing the transverse cross-section of a virion. A cholesterol ester crystal adjacent to the virion is marked by a white arrowhead. **(E)** 3D segmentation of the viral membrane, nucleocapsid and VP40 shown in (D). **(F)** Magnified view of a different slice of the tomogram in (B) showing a longitudinal cross-section through a virion. A linear membrane fragment adjacent to the virion is marked with a white arrow. **(G)** 3D segmentation of the viral membrane, nucleocapsid and VP40 displayed in (F). **(H)** Slices through a tomogram showing a purified Ebola virus before infection. **(I-J)** Magnified areas highlighted in (F) and (H), respectively, showing the viral membrane and VP40 densities at the luminal side. For comparison, line profiles at 3 nm distance from the inner membrane monolayer, visualized by dotted profiles (magenta and blue, respectively), were determined. **(K)** Line profiles adjacent to the inner viral membrane leaflet of a virion inside an endosome and a purified virion before infection. Scale bars: 200 nm (B, H), 50 nm (D, F), 20 nm (I, J).

### Low pH triggers disassembly of the Ebola virus matrix in vitro

We next sought to identify factors driving VP40 disassembly. Since EBOVs enter host cells via late endosomes, which are characterized by low pH, we assessed the effect of external pH on the shape of Ebola virus-like particles (VLPs) and, in particular, on the structure of the VP40 matrix. VLPs composed of VP40 and GP were produced from HEK 293T cells and analyzed by cryo-ET (Fig. 2 A-D). At neutral pH, the organization of VP40 proteins into a helical scaffold was apparent from transverse cross-sections as an additional profile adjacent to the inner membrane monolayer and as regular striations spanning the width of the particles when observed close to the VLP surface (Fig. 2 B, C). Individual VP40 proteins were visible as distinct densities lining the membrane (Fig. 2 D). To understand their organization within the matrix, we applied subtomogram averaging of the VP40 matrix in purified VLPs. In accord with recently published data^28^, the subtomogram average revealed the linear arrangement of VP40 dimers via their C-terminal domains (CTDs), which are directly connected to the inner membrane monolayer (Fig. 1 E). The available crystal structure of the VP40 dimer (pdb: 7jzj) fitted well into the average (Fig. 2 F) except for three short helical segments of one VP40 monomer (Fig. S5 A).

**Figure 2:**
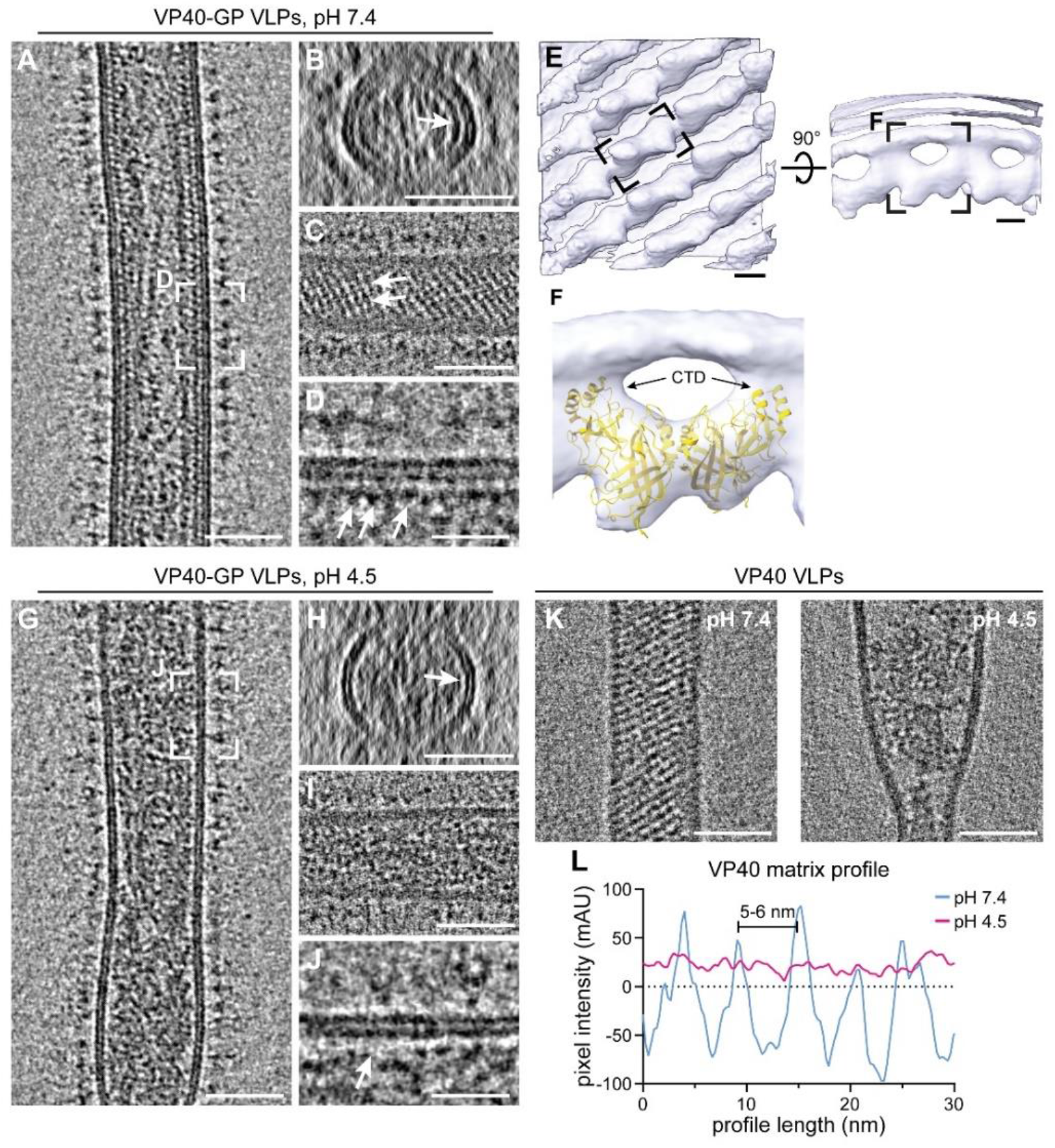
The VP40 matrix in Ebola VLPs disassembles at low pH. **(A)** Slices of a tomogram showing a filamentous Ebola VLP composed of VP40 and GP at neutral pH, (n=37). **(B-C)** Transverse cross-section and longitudinal near-to-surface slices of the tomogram shown in (A) displaying the densities for the outer and inner membrane monolayer and an additional density of the VP40 matrix apparent as striations in (C) (white arrows). **(D)** Longitudinal cross-section highlighting the VP40 densities adjacent to the membrane (white arrows). **(E)** Subtomogram average of the VP40 matrix in Ebola VLPs composed of GP and VP40. A density representing a single VP40 dimer is indicated by a black dashed rectangle. **(F)** Crystal structure of the VP40 dimer (pdb: 7jzj) fitted into the subtomogram average with the C-terminal domains (CTDs) indicated by arrows. **(G-J)** Slices of a tomogram showing a filamentous Ebola VLP composed of VP40 and GP after incubation at low pH (n=18). White arrows in (H) and (J) highlight areas adjacent to the VLP membrane devoid of protein densities in contrast to corresponding slices of VLPs at neutral pH. **(K)** Slices of tomograms showing filamentous VLPs composed of VP40 after incubation at neutral (n=22) and low pH (n=8), respectively. **(L)** Line density profiles determined adjacent to the inner membrane monolayer of VLPs incubated at neutral (blue) and low pH (magenta). At neutral pH, the VP40 matrix detectable as regular densities in (D) have a 5-6 nm pitch. Scale bars: (A-C), (G-I) and (K): 50 nm, (E): 2.5 nm, (D), (J): 20 nm.

To assess whether the VP40 matrix undergoes disassembly at low pH, VLPs were then subjected to the late endosomal pH of 4.5 for 30 min. Consistent with Ebola virions found in late endosomes, the VLPs retained their overall filamentous morphology but did not show ordered VP40 matrix layers. Instead, they contained disordered protein aggregates accumulated at the VLP core (Fig. 2 G-J). Additionally, a lack of densities between the membrane and protein aggregates indicates that VP40 detaches from the membrane, as particularly apparent from the cross-sections (Fig. 2 H, J), which was also reflected in a more variable particle diameter (Fig. S5 B). To elucidate whether this phenotype depends on the presence of EBOV GP, VLPs composed of VP40 alone were analysed by cryo-ET. The presence and absence of the ordered VP40 matrix at neutral and low pH, respectively, were clearly apparent as regular striations and disordered protein accumulations at the particles’ cores (Fig. 2 K, Fig. S5 C). Accordingly, line density profiles proximal to the inner membrane monolayer of VLPs showed the 5-6 nm pitch of the assembled VP40 matrix at neutral pH, whereas no repeating densities were detected at low pH (Fig. 2 L). Hence, pH-mediated VP40 disassembly is independent of other viral proteins.

### VP40 interactions with negatively charged lipids are weakened at low pH

To further probe the specific VP40-lipid interactions at neutral and low pH, we performed all-atom molecular dynamics (MD) simulations and modelled the binding of VP40 dimers to membrane lipids at different pH levels. To this end, we emulated a simplified membrane containing 30% phosphatidylcholine, 40% cholesterol and 30% phosphatidylserine mimicking the overall negative charge of the VLP inner membrane monolayer. We modelled missing C-terminal residues, which are inherently flexible and disordered, into the VP40 dimer structure (pdb: 7jzj) and simulated VP40-membrane interactions for a cumulative time of 10 microseconds for each pH using the CHARMM36m force field^29–31^. We show that after one CTD of the VP40 dimer established interactions with phosphatidylserines, the second CTD is pulled towards the membrane, leading to the anchoring of the dimer into the membrane (Fig. 3 A). The membrane interactions were driven by positively charged residues decorating the C-termini of the VP40 dimer, including K224, K225, K274, and K275, which corroborates experimental data showing that these residues form a basic patch required for membrane association and budding of VLPs^32^. In the MD simulations, the basic patches strongly promote lipid interactions and localize in flexible loops at the CTDs, which penetrate into the inner membrane monolayer (Fig. 3 B, C) and correspond to the previously unassigned densities^8^ between the VP40 matrix and viral membrane in the subtomogram average (Fig. 2 E, F). Moreover, the MD simulations showed that the rotation angle of VP40 monomers oscillates around 1° (SD 9.5) along the N-terminal -dimerization domain and is in agreement with the subtomogram average (Fig. 3 B, Fig. S6 C-E), such that only flexible loops protrude from the average (Fig. 3 D). Accordingly, when aligning the crystal structure of the VP40 dimer (pdb: 7jzj) with the VP40 structure obtained from the MD simulations, the membrane-proximal loops and short alpha-helices were mismatched while the core of the monomer aligned well (Fig. 3 B, highlighted in yellow, Fig. S6 A, B). The second monomer displayed similar secondary structures, which were tilted with respect to the crystal structure by 17°, causing a mismatch when compared to the crystal structure (Fig. 3 B, blue monomer, Fig. S6 E).

**Figure 3:**
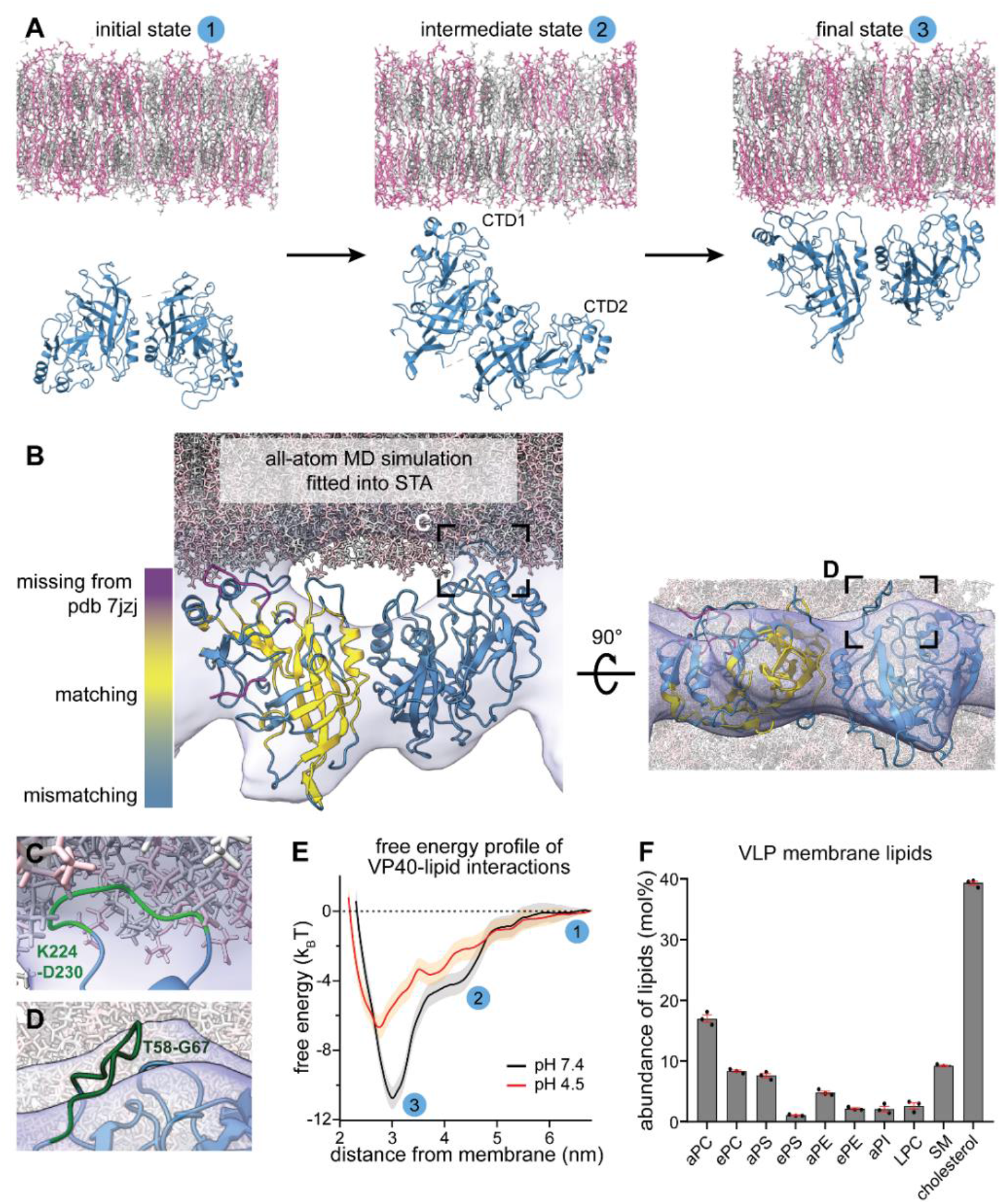
VP40–lipid interactions at neutral and low pH. **(A)** Initial, intermediate, and final state of the VP40-membrane interaction pathway sampled with unbiased all-atom molecular dynamics (MD) simulations. The simulated membrane is composed of 30% phosphatidylcholine, 40% cholesterol, and 30% phosphatidylserine. VP40 is randomly oriented towards the membrane in the initial state. Lipid interactions are first mediated via one C-terminal domain (CTD1) (intermediate state) before the second CTD (CTD2) is ultimately pulled towards to membrane. **(B)** MD simulation frame of the VP40-membrane-bound state, with a rotation angle of VP40 monomers along the N-terminal-dimerization domain (Fig. S2, C-D) of 1°, fitted into the subtomogram average shown in (Fig. 2 E). Missing C-terminal residues in the crystal structure of the VP40 dimer (pdb: 7jzj) were computationally modelled (magenta). VP40 conformational changes upon lipid-interaction resulted in a displacement of secondary structures (steel blue), while the core of the protein remained unaltered in comparison to the crystal structure (yellow). **(C)** The area highlighted in (B) shows a flexible, C-terminal loop (green) containing the residues K224 and K225 that interact with phosphatidylserines in the inner membrane monolayer. **(D)** Area highlighted in the rotated MD simulation in (B) showing a flexible loop (residues T58-G67) protruding from the subtomogram average. **(E)** Free energy profiles of VP40-lipid interactions at pH 7.4 and pH 4.5 determined from MD simulations. The plot shows free energy (in kBT) at increasing membrane-VP40 distances (nm) with indicated three states shown in (A). **(F)** Ebola VLP lipid composition showing highly abundant lipids determined by mass spectrometry in mol%, n=3. Lipid abbreviations: phosphatidylcholine (PC), phosphatidylserine (PS), phosphatidylethanolamine (PE), phosphatidylinositol (PI), lyso-phosphatidylcholine (LPC), sphingomyelin (SM). Prefix “a” indicates acyl-linked glycerophospholipids, prefix “e” indicates ether-linked (plasmanyl) or the presence of one odd and one even chain fatty acyl.

Next, we simulated VP40–membrane interactions at pH 4.5 and observed a significantly decreased affinity towards the membrane, consistent with our tomography data (Fig. 3 E). The free energy profile determined from the MD simulations (Fig. 3 E) revealed an energy minimum that was 4.1 k_B_T weaker at low pH compared to pH 7.4. However, binding was not completely diminished since 10% of the phosphatidylserines used in the simulation are still charged^33^, and the membrane modelled here containing high levels of phosphatidylserine can still engage in electrostatic interactions. To identify which lipids are enriched in the VLP membrane and are thus likely involved in VP40 binding, we then determined the VLP lipid composition by mass spectrometry (Fig. 3 F, Fig. S6, Supplementary table T1). As expected for Ebola VLPs budding from microdomains in the plasma membrane^34,35^, the Ebola VLP envelope was rich in phosphatidylserine and cholesterol, phosphatidylcholine and sphingomyelin (9%, 39%, 25% and 9%, respectively). Collectively, this data argues for low pH-mediated VP40 disassembly through neutralization of negatively charged phospholipids in the viral envelope and highlights electrostatic interactions as the main driving forces of the VP40-membrane binding (Fig. S6 F).

### Protons passively equilibrate across the EBOV membrane

We next assessed the acidification kinetics to elucidate the mechanism of ion permeability across the viral membrane. Ebola VLPs composed of GP, VP40, and the pH-sensitive GFP variant pHluorin^36^ N-terminally fused to VP40 (pHluorin-VP40) were prepared to monitor pH changes in VLP lumina upon altering the pH of the surrounding buffer (Fig. 4 A). Pleomorphic VLPs containing VP40 in excess over VP40-pHluorin, including filamentous and spherical particles, were imaged by time-lapse microscopy (Fig. 4 B, C). At neutral pH, the VLPs showed a fluorescent signal, which gradually decayed over several minutes after lowering the external pH (Fig. 4 D). In contrast, when adding the detergent Triton X-100 (T-X^100^) before imaging to permeabilize the VLP membrane, the signal decayed to background fluorescence within the first 15 seconds (Fig. 4 D), indicating that protonation of pHluorin was slowed down by the membrane of the VLPs. To calculate the acidification kinetics of the VLPs’ lumen, we determined pH levels in the VLPs (Fig. 4 E) by correlating the pHluorin fluorescence intensity to pH using a calibration curve (Fig. S7 A). We found that the luminal pH of filamentous VLPs decreased from 7.4 to 6.4 after 6 minutes, while for the spherical particles, this decay had already occurred after 3.5 minutes (Fig. 4 E). We next calculated the membrane proton permeability coefficient, *P_m_*, based on the geometry of the VLPs measured by cryo-ET (Fig. 2) and the fluorescence decay times (Fig. S7 B). Filamentous VLPs had a permeability coefficient of 1.2 ± 0.2 Å·sec^−1^, whereas the membrane of spherical VLPs was significantly more permeable with a permeability coefficient of 33 ± 9 Å·sec^−1^.

**Figure 4:**
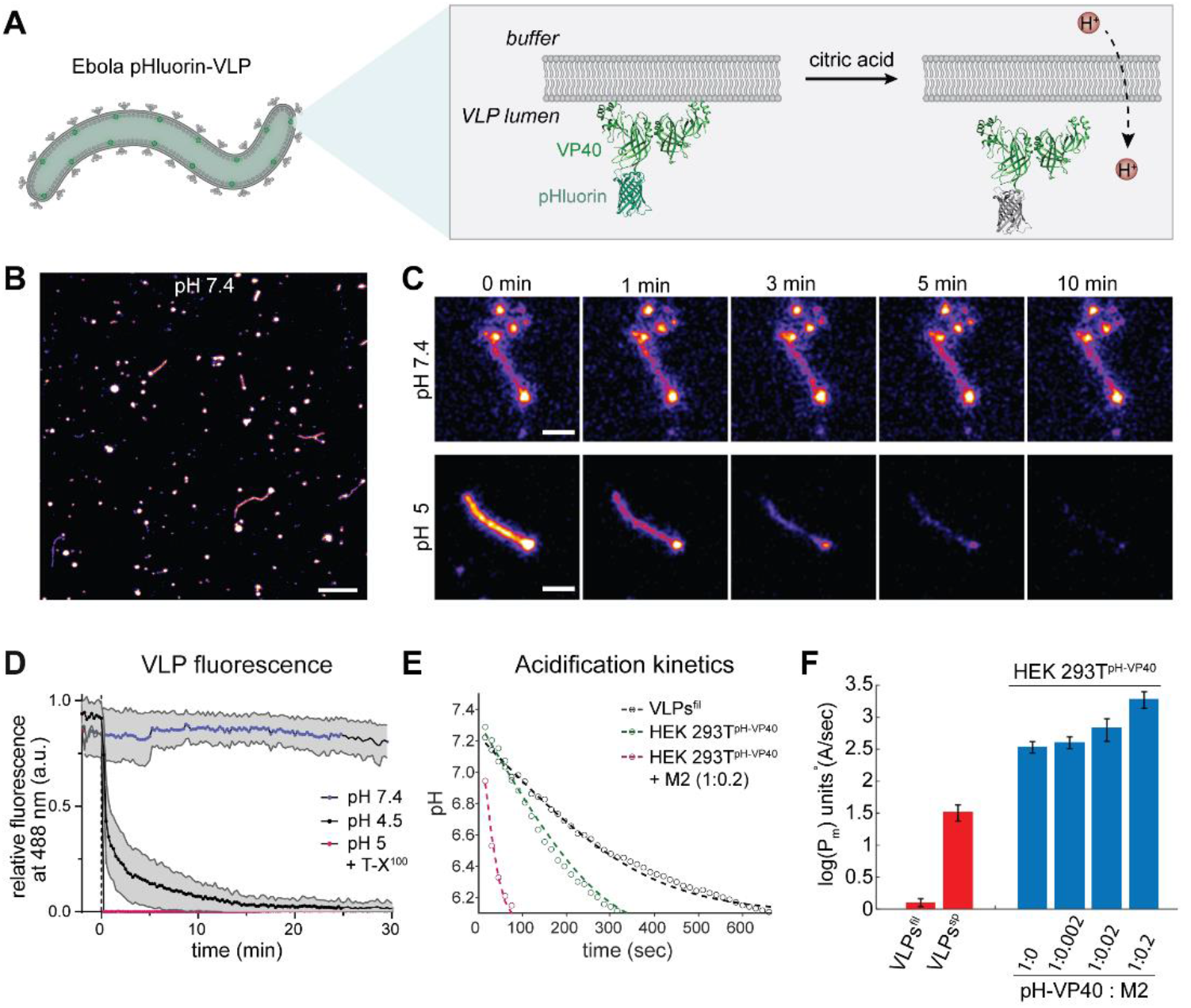
Time-lapse microscopy of Ebola VLPs at different pH. (A) Schematic showing the VLP membrane and pHluorin-VP40 facing the luminal side of the VLPs. Upon protonation, pHluorin loses its fluorescence properties and serves as a proxy for proton diffusion across the membrane. **(B)** Overview confocal fluorescence microscopy image showing pleomorphic pHluorin-labelled VLPs composed of VP40, pHluorin-VP40 (ratio 10:1) and GP. **(C)** Magnified images of representative VLPs acquired during time-lapse microscopy at neutral pH and after acidification to approximately pH 5. Frames are exemplarily shown at 0, 1, 3, 5 and 10 min after lowering the external pH. **(D)** Plot showing the mean relative fluorescence intensities and standard deviation of VLPs imaged at neutral pH, low pH and in the presence of Triton X-100 (T-X^100^) at low pH over time. **(E)** Plot showing the drop of pH inside VLPs over time after lowering the pH of the surrounding buffer to 5. The dots represent the mean values, and the dashed lines are the theoretical fit to Eq. 3. **(F)** Membrane permeability of VLPs (red) and HEK 293T cells expressing different ratios of VP40 and M2 (blue). The permeability is displayed on a logarithmic scale. Permeability coefficients: filamentous VLPs 1.2 ± 0.2 Å ·sec^−1^, spherical VLPs 33 ± 9 Å ·sec^−1^, cells expressing no M2 345 ± 71 Å ·sec^−1^, cells expressing pHluorin-VP40 and M2 at 1:0.002 molar ratio 409 ± 85 Å ·sec^−1^, cells expressing pHluorin-VP40 and M2 at 1:0.02 molar ratio 683 ± 263 Å ·sec^−1^ and cells expressing pHluorin-VP40 and M2 at 1:0.2 molar ratio 1940 ± 562 Å ·sec^−1^.

To compare the membrane permeability of the VLPs with the permeability of membranes containing a well-characterized viral ion channel, we used HEK 293T cells expressing VP40-pHluorin and the influenza ion channel M2. In line with previous measurements^37,38^, the plasma membrane in cells displayed a permeability coefficient of 345 ± 71 Å·sec^−1^ (n=44) in the absence of M2. As expected, the permeability increased with increasing amounts of M2 present in the plasma membrane up to 1940± 562 Å·sec^−1^ (n=26) when M2 and VP40 were transfected at a 1:0.2 molar ratio (Fig. 4 E, F). Compared to the envelope of filamentous Ebola VLPs, the plasma membrane was more permeable to protons already in the absence of M2.

### Disassembly of the VP40 matrix is critical for membrane fusion

Collectively, our experimental data and MD simulations indicate that low pH drives VP40 matrix disassembly and detachment from the viral envelope. We speculated that this influences the GP-mediated membrane fusion between the EBOV envelope and the endosomal membrane. To test this hypothesis, we numerically simulated the membrane fusion pathway in the presence of the VP40 matrix and estimated the magnitude of the two major energy barriers to membrane fusion: stalk and fusion pore formation ^39–42^. We applied a continuum approach to model the lipid membrane with the commonly used framework of the theory of splay-tilt deformations^43,44^ and the VP40 matrix layer as a uniform thin shell that interacts continuously with the virus envelope but can also locally detach from the membrane near the stalk and diaphragm rim (Fig. 5 A). Based on the VP40-membrane binding energy obtained from the MD simulations at pH 7.4 (Fig. 3 G) and the density of VP40 dimers on the viral envelope determined from the subtomogram average (Fig. 2 E), we estimated the VP40 matrix interaction energy density to be 0.38 ± 0.02 k_B_T/nm^2^ (with a dimer density of 0.036 nm^−2^ and free binding energy of 10.77±0.47 k_B_T). Consistently with our cryo-ET data, we assume that the interaction energy density vanishes at low pH due to the VP40 matrix disassembly. Importantly, our calculations showed that the initiation of viral membrane fusion is more favorable after VP40 disassembly. The calculated stalk formation energy barrier drops from 89-79 k_B_T to 65 k_B_T due to the weakening of the VP40-lipid interactions at low pH, depending on the matrix layer rigidity (Fig. 5 B). Hence, the intact VP40 matrix can prevent or slow down the stalk formation, which is the first step of membrane fusion.

**Figure 5:**
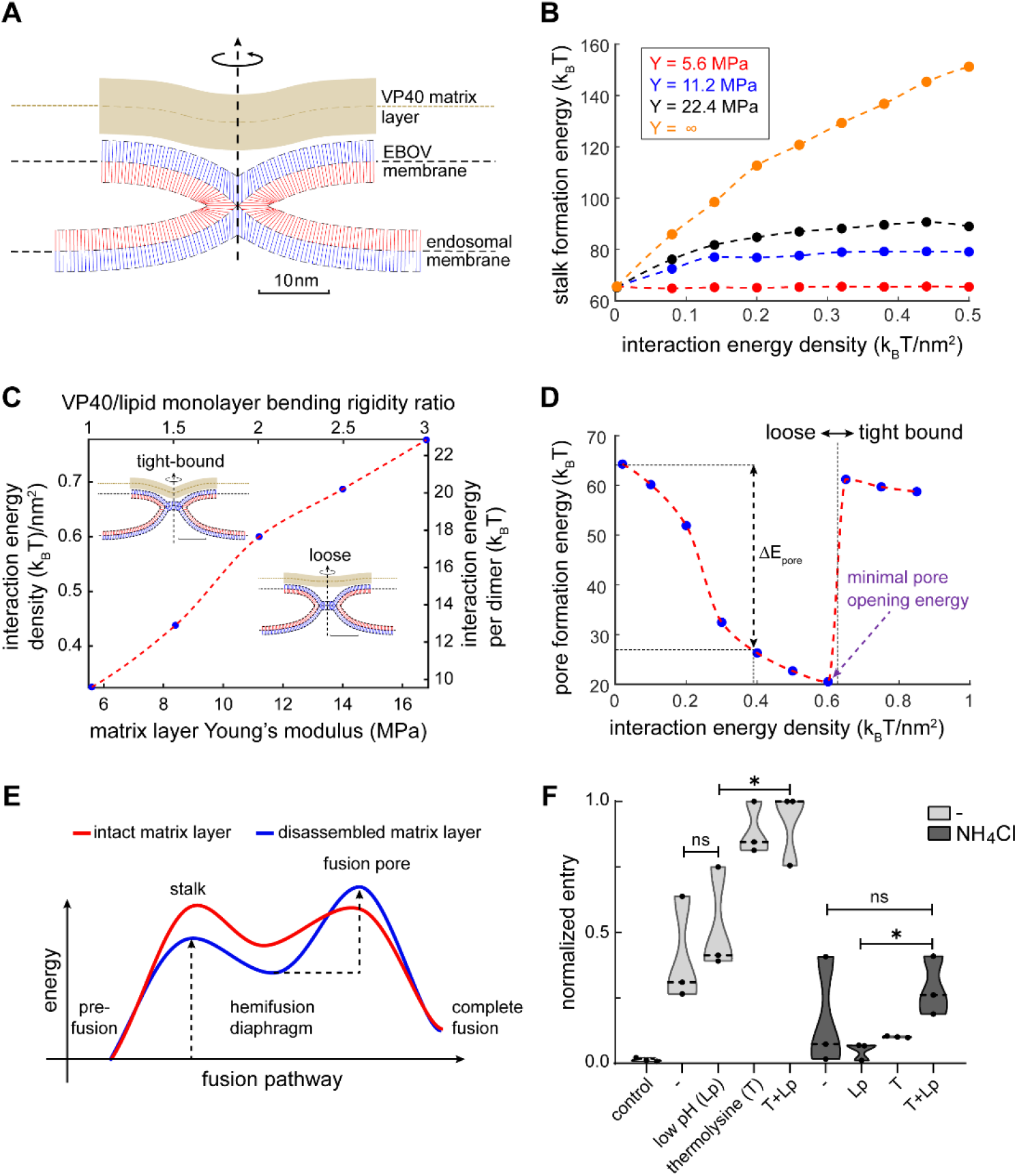
Membrane fusion dynamics in the presence and absence of the VP40 layer. **(A)** Simulation result of a hemifusion stalk in the presence of a rigid matrix layer (VP40). The blue and red lines represent the averaged lipid director of the distal and proximal monolayers, respectively. The VP40 matrix layer is represented by the continuous thick brown strip. Parameters used in panels (A-D) for the lipid membrane: lipid monolayer bending rigidity 10 k_B_T, tilt decay length 1.5 nm, saddle splay modulus to bending modulus ratio −5 k_B_T, monolayer spontaneous curvature −0.22 nm^−1,^ and monolayer width 1.5 nm. VP40 matrix layer: width 4 nm, Poisson’s ratio 0.5, and membrane mid-plane to VP40 mid-plane optimal distance 4 nm. In panel (A) the matrix layer Young’s modulus is 11.2 MPa, and the interaction energy density is 0.2 k_B_T/nm^2^. **(B)** Stalk formation energy as a function of VP40-membrane interaction energy. The stalk energy for non-interacting VP40 matrix (*u*_0_ = 0) is 65 k_B_T. VP40 matrix layer Young’s modulus legend – red 5.6 MPa, blue 11.2 MPa, black 16.8 MPa, and orange is infinitely rigid. The bending rigidity ratio between the VP40 matrix layer and lipid monolayer are 1, 2, 3, and infinity, respectively. The line represents an infinitely rigid layer. **(C)** Hemifusion diaphragm configurations phase-diagram – above dotted red line: tight-bound solution and loose configuration below. The inserts are simulation results with layer Young’s modulus of 11.2 MPa. The interaction energy density is 0.2 k_B_T/nm^2^ in loose configuration and 0.85 k_B_T/nm^2^ in the tight-bound configuration. The scale bar is 10 nm. **(D)** Fusion pore formation energy as a function of VP40-membrane interaction energy. The discontinuity in the energy is located at the phase line between configurations (see C). The change in pore formation energy, Δ*E*_*pore*_ is defined as the difference between fusion-pore formation energy at interaction energy density 0.2 k_B_T/nm^2^ (the value found using MD simulations) to the matrix-free case (no interaction energy). A-D) Dotted lines serve as a guide to the eye. **(E)** Illustration of the effect of the matrix layer on the fusion pathway and the fusion intermediates in the absence of the matrix layer. As a result of the presence of the matrix layer, the stalk formation energy barrier increases while the pore formation energy barrier decreases and the hemifusion diaphragm intermediate is less stable. **(F)** Quantification of FACS data showing Ebola VLP entry as measured by a fluorescence shift of infected cells from emission at 510 nm (no entry) to 450 nm (entry). VLPs were treated prior to infection as indicated on the x-axis, with control: uninfected control cells, – : no treatment, T: thermolysin-treatment at neutral pH, Lp: low pH treatment. Target cells were treated with media or ammonium chloride (NH_4_Cl), n= 3 with 10’000 cells measured per sample.

Interestingly, our model predicts that fusion pore formation, which occurs after stalk formation, is facilitated in the presence of the assembled VP40 matrix because of increased stress in the hemifusion diaphragm. Simulation data showed that the interaction energy density and the rigidity of the VP40 matrix modulate the shape of the hemifusion diaphragm structure (Fig. 5 C), which determines the energy barrier of pore formation. Strong interactions between the lipids and VP40 matrix (Fig. 5 C, ‘tight-bound’ configuration) stabilize the hemifusion diaphragm, thereby inhibiting fusion pore formation. Conversely, in case of a weakly interacting or stiff VP40 matrix (Fig. 5 C, ‘loose’ configuration), the hemifusion diaphragm is more unstable, which results in a lower energy barrier for fusion pore formation (Fig. 5 D). Our model showed that the minimal pore opening energy is at the phase boundary between ‘loose’ and ‘tight-bound’ configurations, where diaphragm stress is maximal (Fig. 5 D). Given the VP40-membrane binding energy and VP40 dimer envelope density found in the MD simulations (Fig. 3 E), we could show that the VP40 matrix and the membrane preferably adopt the ’loose’ configuration at both neutral and acidic pH. Therefore, contrary to the stalk formation energy barrier, which is decreased upon VP40 matrix disassembly, the pore formation energy barrier is lower in the presence of the VP40 matrix layer by 16-33 k_B_T (Fig. 5 D), depending on the matrix layer rigidity (Fig. S4 A).

To validate this theoretical model experimentally, we performed beta-lactamase entry assays using VLPs^45^. EBOV membrane fusion requires proteolytic cleavage of GP by low pH-activated cathepsin proteases^16^ and subsequent binding of the cleaved GP1 subunit to the endosomal receptor NPC1. To circumvent the need for low pH to activate cathepsin proteases, we substituted cathepsins with thermolysins which are active at neutral pH^46^, thereby decoupling the low pH requirement from proteolytic GP processing. Ebola VLPs composed of GP, VP40, and beta-lactamase N-terminally fused to VP40 (BlaM-VP40) were purified and subjected to thermolysin treatment followed by incubation at neutral or low pH. We then incubated target Huh7 cells with the pre-treated VLPs, loaded the cells with a fluorescent BlaM substrate, and assessed virus entry by FACS (Fig. S8). Thermolysin treatment significantly enhanced host cell entry, whereas the enhancement of entry by low pH treatment alone was less pronounced and not statistically significant (Fig. 5 F). To determine whether thermolysin-treated VLPs still require low pH for entry, we challenged host cells treated with ammonium chloride, which blocks endosomal acidification. Strikingly, entry of VLPs treated with thermolysin was completely inhibited by ammonium chloride, which is in line with a previous study conducted with bafilomycin to inhibit endosomal acidification^47^. This suggests that GP processing alone is insufficient to enable entry. Conversely, low pH treated VLPs were also unable to enter target cells treated with ammonium chloride since impaired endosomal acidification prevents the activation of cathepsin proteases and hence GP priming. Combined thermolysin- and low pH-treatment of VLPs *in vitro* rescued entry into host cells with inhibited endosomal acidification, albeit to a lesser extent compared to entry into untreated cells. Overall, these data show that VP40 matrix integrity modulates GP-mediated membrane fusion, strongly supporting the notion that VP40 disassembly is required for and precedes membrane fusion.

## Discussion

EBOVs are remarkably long, filamentous virions that enter the cytoplasm by fusion with late endosomal membranes. Similar to other enveloped viruses, the shape and stability of EBOVs are determined by a matrix layer forming a flexible scaffold underneath the viral envelope, which is indispensable for particle formation and protects the encapsidated genome during transmission. Here, we investigate the molecular architecture of the EBOV VP40 matrix in Ebola virions during host cell entry to elucidate whether and how it is released from the viral envelope to allow virion uncoating. Using *in situ* cryo-electron tomography, we directly visualize EBOVs entering host cells via the endosomal route. Virions inside endosomal compartments exclusively exhibited disassembled VP40 matrices and some had engulfed endosomal vesicles, suggesting that the membranes of these virions are sufficiently flexible to engage in membrane fusion (Fig. 1). Considering that the nucleocapsids in all endosomal EBOVs were condensed, we propose that VP40 disassembly precedes membrane fusion, while nucleocapsid integrity is maintained until cytoplasmic entry is concluded. The VP40 aggregation surrounding the nucleocapsid may be involved in engaging cellular factors required to pull nucleocapsids out of the fusion site as has recently been suggested for influenza A virus, whose disassembled M1 matrix layer recruits the aggresome machinery by mimicking misfolded proteins^21^. Supported by our functional data and computational simulations, we propose that EBOV uncoating occurs in a cascade-like fashion. Tightly regulated by pH, uncoating starts with the disassembly of the VP40 layer, followed by GP-driven membrane fusion and release of the compact nucleocapsid into the cytoplasm. It remains to be elucidated when and how the nucleocapsid undergoes de-condensation to allow viral genome replication and transcription.

The organization of VP40 proteins within the VP40 matrix, including their oligomeric state and orientation of C-termini towards the membrane, has long been subject of debate^34,48,52–55^. While the structure of VP40 in solution was revealed as a dimer^32^, structures of VP40 in the context of lipid environments were proposed based on purified VP40 proteins either truncated or characterized in the presence of lipid mimics. These revealed VP40 hexamers as the building blocks of theVP40 matrix^32,56,57^, in which the C-termini alternatingly face the viral membrane. Recently published data^28^ and our subtomogram averaging (Fig. 2) show that the VP40 matrix within VLPs is instead composed of linearly arranged dimers, in which all C-termini are facing the VLP membrane and thus collectively contribute to the electrostatic interactions. Importantly, a combination of MD simulations and subtomogram averaging allowed us to refine the structure of the VP40 dimer interacting with the membrane and to map the basic patch of lysine residues to flexible loops that extend into the inner membrane monolayer^58^ (Fig. 3). Additionally, our MD simulations reveal lipid-induced conformational changes of the VP40 dimer that complement our subtomogram averaging data. The rotation of VP40 monomers along the N-terminal-dimerization domain is in line with the structural data and emphasizes the modularity of the VP40 dimer, which may contribute to the flexibility of the large filamentous particles^8,59^.

Using VLPs of minimal protein composition (VP40 and GP, and VP40 alone), we show that VP40-disassembly, i.e. the detachment of the matrix from the viral envelope is triggered by low endosomal pH (Fig. 2). This indicates that VP40 disassembly does not depend on structural changes of other viral proteins and is driven solely by the acidic environment. Furthermore, we deduced VP40-lipid interaction strengths from the MD simulations, which are strongly diminished at pH 4.5 and thus support a dissociation of VP40 from the membrane in endosomal environments. Our data demonstrate that VP40 detachment from the membrane is driven by the neutralization of negatively charged phospholipids at endosomal pH. VP40 detachment from viral envelope is caused by a disruption of electrostatic interactions between VP40 and negatively charged lipids in the viral envelope, which have experimentally been demonstrated and attributed to a basic patch of lysine residues decorating the VP40 C-termini^48–51^. Considering that matrix protein assembly of other RNA viruses relies on electrostatic interactions with negatively charged lipids^60^, we propose that pH-mediated matrix disassembly is a general mechanism critical for viral uncoating.

Notably, pH-driven structural remodeling of viruses has so far only been shown and extensively studied for influenza A virus^61^, which is known to encode the viral ion channel M2 (reviewed here^62^). Since EBOVs do not encode a dedicated ion channel, we determined the permeability of the Ebola VLP membrane in comparison to the plasma membrane in the absence and presence of the M2 ion channel. We show that the proton permeability of the VLP membrane depends on particle morphology and is markedly lower in filamentous VLPs when compared to spherical VLPs (Fig. 4). Since spherical Ebola virions are predominantly released at very late infection time-points (4 days post infection) and are less infectious than filamentous particles^63^, it is plausible that their membrane properties including proton permeability result from improper particle formation due to cell exhaustion. The higher proton permeability of the plasma membrane already in the absence of M2 likely results from its complex composition comprising host cell ion channels^64^. While the membrane permeability of filamentous VLPs is low compared to values reported in the literature for protein-free liposomes^37^, pH equilibration inside filamentous virions is fast due to their small radius and takes place within minutes. This suggests that acidification occurs rapidly after EBOV uptake into late endosomes and is not rate-limiting during virus entry into host cells, in agreement with a previous report^65^. Taken together, we show that protons diffuse passively across the EBOV envelope, independent of a dedicated ion channel. It remains to be elucidated whether virion acidification also occurs by passive diffusion in other late-penetrating viruses lacking a dedicated ion channel.

We further show that the energy barriers of both the hemifusion stalk and fusion pore formation strongly depend on the VP40 matrix rigidity (Fig. 5). The assembled VP40 matrix inhibits stalk formation, which precedes fusion pore formation during membrane fusion, arguing for VP40 disassembly as a critical step required for membrane fusion and highlighting the role of the matrix as a modulator of membrane fusion. Together, the findings presented here reveal a yet unknown role of viral matrix proteins during viral entry and uncoating as membrane fusion modulators. We propose that low-pH driven matrix protein disassembly is decisive for membrane fusion of other enveloped late-penetrating viruses, making the process a promising target for interventions by development of virus matrixspecific weak base inhibitors.

## Materials and Methods

### Cell lines and Ebola VLP production

Cell lines used in this work include HEK 293T cells for Ebola virus-like particle (VLP) production, and Huh7 cells as target cells to assess VLP and EBOV entry. HEK 293T were obtained from ECACC General Cell Collection and Huh7 cells were kindly provided by Prof. Ralf Bartenschlager (Heidelberg University Hospital). Both cell lines were maintained in DMEM media (ThermoFisher Scientific) supplemented with 10% (v/v) FBS and 100 U/ml penicillin-streptomycin (ThermoFisher Scientific) at 37°C, 5% CO_2_. All cells were tested for Mycoplasma contamination every 3 months. Ebola VLPs were produced by transfecting HEK 293T cells with equal amounts of pCAGGS plasmids encoding EBOV GP, VP40, NP, VP35 and VP24 (species *Zaire ebolavirus*, Mayinga strain). Supernatants of transfected cells were harvested 48 h post transfection and clarified by centrifugation at 398 × g for 10 min, and 2168 × g for 15 min (JA-10 rotor, Beckmann). Clarified supernatants were passed through a 30 % sucrose cushion in HNE buffer (10 mM HEPES, 100 mM NaCl, 1 mM EDTA, pH 7.4) by centrifugation for 2.5 h at 11,400 rpm (SW32 Ti rotor, Optima L-90K ultracentrifuge, Beckmann). Pellets were resuspended in HNE buffer and centrifuged at 19,000 rpm (TLA 120.2 rotor, Optima TLX ultracentrifuge (Beckmann)) to remove residual media and sucrose. Final pellets were resuspended in HNE buffer and protein concentrations were measured using the Pierce BCA Protein Assay Kit (ThermoFisher Scientific) according to the manual provided by the manufacturer.

To produce reporter VLPs, pHluorin was N-terminally cloned to VP40 and beta-lactamase-VP40 (BlaM-VP40) was a kind gift from Dr. Kartik Chandran. Reporter VLPs were produced by transfecting EBOV GP, VP40, and pHluorin-VP40 or BlaM-VP40 in a 10:10:1 ratio and purified as described above.

### Production of Ebola virus and infection of Huh7 cells

EBOV (species *Zaire ebolavirus*, strain Mayinga) was produced in VeroE6 cells in the BSL4 facility at the Friedrich-Loeffler Institut (Insel Riems, Greifswald), following approved standard operating procedures. 5 days post-infection, supernatants of infected cells were harvested and purified as described for the VLPs above, and then fixed by adding paraformaldehyde and glutaraldehyde in HNE buffer for a final concentration of 4% and 0.1%, respectively.

For structural characterization of EBOV-infected cells, Huh7 cells were seeded on 200 mesh Quantifoil™ SiO_2_ R1.2/20 EM grids placed on 3D-printed grid holders in a 96-well plate. 0.0075 × 10^6^ cells were seeded, and the plates were transferred to the BSL4 laboratory after 4-5 h. Cells were infected with unpurified EBOVs at an MOI of 0.1 and incubated for 48 h before chemical fixation for 24 h with 4 % paraformaldehyde and 0.1 % glutaraldehyde in PHEM buffer (60 mM PIPES, 25 mM HEPES, 2 mM MgCl_2_, 10 mM EGTA, pH 6.9). After transfer of the samples out of BSL4, the grids were kept in PHEM buffer and plunge-frozen within three days.

### Sample preparation for cryo-electron tomography

Ebola VLPs and chemically fixed EBOV were plunge-frozen as previously described^66^. Briefly, VLPs were diluted to approximately 10-20 ng/μl, mixed with 10 nm protein A-coated colloidal gold (Aurion) and applied onto a glow-discharged EM grid (200 mesh, R 2/1, Quantifoil) prior to plunge-freezing with a Leica EM GP2 automatic plunge-freezer.

Chemically fixed EBOV-infected Huh7 cells on EM grids were vitrified using the GP2 plunge-freezer (Leica) at a ethane temperature of −183°C, chamber temperature of 25°C and 95% humidity. 5 μl PHEM buffer were added to the girds before blotting them from the back with a Whatman Type 1 paper for 3 sec. For cryo-focused-ion beam (FIB) milling, the grids were clipped into specifically designed AutoGrids™ (ThermoFisher Scientific).

Cryo-FIB milling was performed as previously described^25^ using an Aquilos dual-beam FIB-SEM microscope (ThermoFisher Scientific). Briefly, cells were selected for milling and coated with an organometallic platinum layer for 5 sec before milling in four successive steps using a gallium-ion beam at acceleration voltage 30 eV. Resulting lamellae were 200-250 nm thick.

### Tomogram acquisition, reconstruction, and volume rendering

Cryo-ET of VLPs and lamellae of EBOV-infected Huh7 cells was performed as previously described^67^. Briefly, data were collected on a Titan Krios Transmission Electron Microscope (TEM, ThermoFisher Scientific) at Heidelberg University operated at 300 keV and equipped with a BioQuantum® LS energy filter with a slit width of 20 eV and K3 direct electron detector (Gatan). Tilt series were acquired at 33,000 ’ magnification (pixel size 2.671 Å) using a dose-symmetric acquisition scheme^68^ with an electron dose of approximately 3 e^−^/Å^2^ per projection with tilt ranges from 60° to −60° in 3° increments using SerialEM (Mastronarde, 2005) and a scripted dose-symmetric tilt-scheme (Hagen et al., 2017).

For subtomogram averaging, tomograms were acquired at EMBL Heidelberg using a Titan Krios TEM (ThermoFisher Scientific) operated at 300 keV and equipped with a Gatan Quantum 967 LS energy filter with a slit width of 20 eV and a Gatan K2xp detector. Tilt series were acquired at 81,000 ’ magnification (pixel size 1.7005 Å) at a defocus range of −3 to −1.5 μm using SerialEM (Mastronarde, 2005) and a scripted dose-symmetric tilt-scheme (Hagen et al., 2017) from −60° to 60° with 3° steps.

Tomograms were reconstructed using the IMOD software package^69^. Stacks of tomograms of VLPs were aligned using gold fiducials, and stacks of tomograms acquired on lamellae were aligned using patch tracking. After 3D contrast transfer function (CTF) correction and dose-filtration implanted in IMOD, the reconstruction was performed by weighted back-projections with a simultaneous iterative reconstruction technique (SIRT)-like filter equivalent to 10 iterations. Tomograms used for subtomogram averaging were reconstructed using 2D CTF correction by phase-flipping and weighted back-projection without a SIRT-like filter. For visualization, 10 slices of the final tomogram were averaged and low-pass filtered.

3D segmentation was performed using the Amira software and the implemented Membrane Enhancement Filter. Membranes were automatically segmented using the Top-hat tool and final segmentations were manually refined.

### Subtomogram averaging

Subtomogram averaging of the VP40 matrix was performed using the Dynamo software package^70,71^. Particles were automatically picked using the filament model and subtomograms were extracted with a cubic side length of 128 voxels from 23 tomograms. A reference template was obtained by iteratively aligning and averaging of 50 subtomograms using a mask permitting alignments only a membrane VP40 layer. The initial average was then used as a template for the final averaging of approximately 7,800 particles.

### Molecular Dynamics Simulations

We used the truncated (residues 45-311) crystallographic structure of the VP40 dimer deposited by Norris, M.J. et al. (pdb: 7JZJ^28^) for atomistic molecular dynamics simulations. The missing CTD loops were modeled using the GalxyFill software^72^ within the CHARMM-GUI web server^73^. The protonation states of the proteins at pH 7.4 and 4.5 were calculated through the PROPKA web server^74^, which indicated a change in the protonation state at pH 4.5 for the following residues: E76, E325, H61, H124, H210, H269, H315. First, the proteins were simulated in water with a 0.1 M NaCl for 1 microsecond. Next, the final structures were placed at a distance of 2 nm from a previously built model membrane surface containing POPC:POPS:CHOL (30:30:40) at ten different random orientations. The model membrane was made using the CHARMM-GUI membrane builder^75^. Since the percentage of POPS charged molecules at pH 4.5 is 10%^33^, we modelled the membrane at pH 4.5 by randomly replacing 90% of POPS molecules with its protonated model (POPSH). Then, each of the ten repeats was solvated with 40913 water molecules and 0.1 M NaCl. Next, charges were neutralized by adding or removing the needed amount of Na^+^- or CL^−^-ions. Finally, each system was simulated for 1 microsecond under NpT conditions. Four out of ten simulations, at both pH conditions, showed VP40 dimer binding to the membrane with the experimentally known binding residues, K224, K225, K274 and K275. These simulations were used for the analysis. For the production run, we employed the Parrinello-Rahman barostat^76^ with a semi-isotropic pressure coupling scheme and a time constant set to 5.0 ps to maintain the pressure constant. The pressure was set to 1.0 bar and the isothermal compressibility to 4.5 × 10–5 bar–1. The temperature was maintained at 310 K using the Nose-Hoover thermostat^77^ with a time constant of 1.0 ps. Electrostatic interactions were handled using the PME method^78^. The cut-off length of 1.2 nm was used for electrostatic (real space component) and van der Waals interactions. Hydrogen bonds were constrained using the LINCS algorithm^79^. Finally, periodic boundary conditions were applied in all directions. The simulations were carried out using an integration time step of 2 fs with coordinates saved every 100 ps. All simulations have been carried out with the GROMACS-2021 software^80^. Protein, lipids, and salt ions were described using the CHARMM36m force field^29–31^. For water, we used the TIP3 model^81^. All pictures, snapshots, and movies were rendered with the Visual Molecular Dynamics (VMD) software^82^.

### Free Energy Calculation

The potential of mean force (PMF) for the VP40 dimer binding on a model membrane surface was calculated using an atomistic resolution, employing the umbrella sampling protocol^83,84^. The initial configuration for each umbrella window was taken directly from unbiased MD simulations. The centre of the mass distance between the VP40 dimer and the phosphate atoms of one leaflet was used as the reaction coordinate. A total of 49 windows, 0.1 nm spaced, were generated and simulated with a harmonic restraint force constant of 2000 kJ·mol^−1^·nm^−2^ for 200 nanoseconds. The first 100 ns of the simulations were considered as an equilibration phase and discarded from the actual free energy calculation. The free energy profiles were reconstructed using the Weighted Histogram Analysis Method^85^. The statistical error was estimated with 200 bootstrap analyses.

### Dihedral angle calculation

The rotation angle of VP40 monomers along the NTD-dimerization domain is defined as the angle between the plane containing the vector connecting alpha carbon atoms of L75^monomer1^ and T112^monomer1^ and the vector connecting atoms T112^monomer1^ and T112^monomer2^ and the plane containing this second vector and the vector connecting atoms T112^monomer2^ and L75^monomer2^ as explained in Fig. S3, A. The angle has been calculated rerunning the simulations trajectory with a GROMACS version patched with the open-source, community-developed PLUMED library^86^, version 2.4^87^. The angle measurement in water has been calculated using all simulation frames. For the angle calculation upon the binding to the membrane, the last 100ns of the four simulations showing VP40-membrane interaction via the experimentally known critical residues (i.e., K224, K225, K274, and K275), have been used.

Regardless of pH, VP40 monomers within the dimer are flexible with a rotation angle, defined as the torsional angle around the alpha carbons of residue T112 of the two monomers (Fig. S2), oscillating around 1° (SD 9.5) in water, which is 17° smaller of the one measured for the crystallographic structure (pdb: 7jzj; Fig. S2). VP40 dimer flexibility is not constrained upon binding to the membrane, however, after binding to the bilayer, the angle distribution was significantly (p ≤ 0.0001) shifted to a value of 3.7° (SD 8.1) and 4.5° (SD 10.7) at pH 7.4 and 4.5 respectively.

### Lipidomics of Ebola VLPs

Ebola VLPs composed of GP, VP40 and the nucleocapsid proteins NP, VP24 and VP35 were produced from HEK 293T cells and purified as described above. They were used at a final protein concentration of 880 ng/μl for lipidomics analysis. VLPs were subjected to lipid extractions using an acidic liquid-liquid extraction method^88^ as described in Malek et al., 2021^89^. In order to ensure that similar amounts of lipids were extracted, a test extraction was performed to determine the concentration of PC as a bulk membrane lipid. Quantification was achieved by adding 1-3 internal lipid standards for each lipid class, with the standards resembling the structure of the endogenous lipid species. Of note, sample volumes were adjusted to ensure that all lipid standard to lipid species ratios were in a linear range of quantification. Typically, the standard to species ratios were within a range of >0.1 to <10. Following this approach, a relative quantification of lipid species was performed. Lipid standards were added prior to extractions, using a master mix consisting of 50 pmol phosphatidylcholine (PC, 13:0/13:0, 14:0/14:0, 20:0/20:0; 21:0/21:0, Avanti Polar Lipids), 50 pmol sphingomyelin (SM, d18:1 with N-acylated 13:0, 17:0, 25:0, semi-synthesized^90^, 100 pmol deuterated cholesterol (D7-cholesterol, Cambridge Isotope Laboratory), 30 pmol phosphatidylinositol (PI, 17:0/ 20:4, Avanti Polar Lipids), 25 pmol phosphatidylethanolamine (PE) and 25 pmol phosphatidylserine (PS) (both 14:1/14:1, 20:1/20:1, 22:1/22:1, semi-synthesized^90^, 25 pmol diacylglycerol (DAG, 17:0/17:0, Larodan), 25 pmol cholesteryl ester (CE, 9:0, 19:0, 24:1, Sigma), and 24 pmol triacylglycerol (TAG, LM-6000/D5-17:0,17:1,17:1, Avanti Polar Lipids), 5 pmol ceramide (Cer, d18:1 with N-acylated 14:0, 17:0, 25:0, semi-synthesized^90^ or Cer d18:1/18:0-D3, Matreya) and 5 pmol glucosylceramide (HexCer) (d18:1 with N-acylated 14:0, 19:0, 27:0, semi-synthesized or GlcCer d18:1/17:0, Avanti Polar Lipids), 5 pmol lactosylceramide (Hex2Cer, d18:1 with N-acylated C17 fatty acid), 10 pmol phosphatidic acid (PA, 17:0/20:4, Avanti Polar Lipids), 10 pmol phosphatidylglycerol (PG, 14:1/14:1, 20:1/20:1, 22:1/22:1, semi-synthesized^90^ and 5 pmol lysophosphatidylcholine (LPC, 17:1, Avanti Polar Lipids). The final CHCl_3_ phase was evaporated under a gentle stream of nitrogen at 37 °C. Samples were either directly subjected to mass spectrometric analysis, or were stored at −20 °C prior to analysis, which was typically done within 1-2 days after extraction. Lipid extracts were resuspended in 10 mM ammonium acetate in 60 μl methanol. Two μl aliquots of the resuspended lipids were diluted 1:10 in 10 mM ammonium acetate in methanol in 96-well plates (Eppendorf twin tec 96) prior to measurement. For cholesterol determinations, the remaining lipid extract was again evaporated and subjected to acetylation as previously described^91^. Samples were analysed on an QTRAP 6500+ mass spectrometer (Sciex) with chip-based (HD-D ESI Chip, Advion Biosciences) electrospray infusion and ionization via a Triversa Nanomate (Advion Biosciences). MS settings and scan procedures are listed in Supplementary table T2. Data evaluation was done using LipidView (Sciex) and an in-house-developed software (ShinyLipids).

### Calibration of pHluorin fluorescence

HEK 293T cells were reverse transfected and seeded at a seeding density of 0.02 × 10^6^ cells per well in a 96 well plate. Briefly, transfection mixtures were prepared containing a pCAGGS plasmid encoding pHluorin-VP40. After 15 min incubation at RT, HEK 293T cells were trypsinized and mixed with the transfection complexes before seeding on a fibronectin-coated 96 well plate. HNE buffers (10 mM HEPES, 100 mM NaCl, 1 mM EDTA) were prepared and calibrated to a pH of 3, 3.5, 4, 4.5, 5, 5.5, 6, 6.5, 7, 7.34, 7.52 and 8. Approximately 20 h post seeding, media was removed, the cells were washed once with HNE buffer at pH 7.34, and incubated for 45 min at 37°C, 5% CO_2_, in the different HNE buffers. Fluorescence intensities were measured at 488 nm excitation using a Tecan plate-reader. To calibrate the fluorescence of pHluorin at different pH, fluorescence was plotted against the pH.

### Permeability of the HEK 293T cell plasma membrane

HEK 293T cells were reverse transfected as described above using pCAGGS plasmids encoding pHluorin-VP40 and influenza M2 at a molar ratio of 1:0, 1:0.0002, 1:0.002 and 1:0.2. Approximately 20 h post seeding, the media was removed, and cells were washed once with HNE buffer, pH 7.34. The buffer was then exchanged with HNE buffer calibrated to pH 4.5 and fluorescence was immediately measured in 15 s intervals using a Tecan plate-reader.

### Time-lapse microscopy of Ebola VLPs

Time-lapse microscopy was performed using a Leica SP8 confocal microscope with a 63 × oil immersion objective. Purified pHluorin-labelled VLPs were added to a glow-discharged μ-Slide 8 well dish (ibidi) at a protein concentration of 10 ng/μl and were allowed to settle for 20 min at RT. They were then imaged using a 488 nm excitation laser and emission at 500-600 nm. Z-stacks were acquired in 15 s intervals for 30 min. To assess acidification kinetics, citric acid was added at a final concentration of 2.6 mM two min after starting the data acquisition. To assess acidification kinetics in the absence of the viral membrane, VLPs were incubated for 5 min with TritonX-100 at a final concentration of 0.1% before imaging.

#### Membrane permeability theory

We estimate the membrane permeability based on the geometry and pH equilibrium time of the VLPs. The membrane total proton flux, *I*, is proportional to the area of the VLPs, *A_VLP_*, ion concentration difference between the buffer, *C_B_*, and VLPs, *C_VLP_*, Δ*C*(*t*) = *C_B_* − *C_VLP_*(*t*), and membrane permeability coefficient *P_m_*^37^,

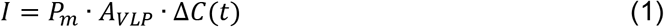

It is easy to show that the protons concentration difference decays exponentially with time t,

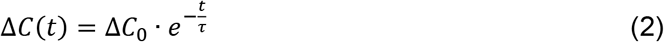

With the decay time constant 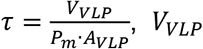 the volume of the VLPs and Δ*C*_0_ the initial concentration difference. The pH level is the logarithm of the protons concentration and can be related to the concentration difference as follows:

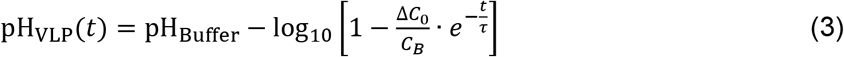

Next, we use a least-squares minimization procedure to fit the measured pH to Eq. 3. We find the three minimization parameters pH_Buffer_, 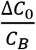 and *τ*. Since the VLPs are either spherical or filamentous, we can derive the membrane permeability coefficient 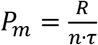, with *R* the respective radius and *n* is either 2 for filamtoues VLPs or 3 for spherical VLPs and cells. The fitted decay times *τ* are presented in Fig. S3 B and the VLPs radii are found using cryo-ET (Fig. S3). In line with previous measurements of Ebola VLPs and virions^8^, the filamentous VLPs had an average radius of 34 ± 4.5 nm (n = 90), while spherical particles are more heterogeneous in size, with a radius of 426 ± 100 nm (n = 12). A similar analysis was also performed on HEK 293T cells. The cells had a round shape. The radius was estimated using fluorescence microscopy to be 17.5 ± 2.5 nm.

### Membrane fusion in the presence of a matrix layer

The fusion process involves three players – the membrane, the matrix layer, and the fusion proteins. In the following section, we describe the physical properties of these three, their interaction, and the fusion pathway in the presence of a matrix layer. We determine the effect of the matrix layer on fusion rate by calculating the magnitude of the two major energy barriers to membrane fusion ^41,42,92^ – stalk formation and fusion pore expansion in the presence of the matrix layer and compare it to the matrix-free state.

#### Description of the fusion site and the fusion process

The fusion reaction starts when the fusion proteins bring the EBOV and endosomal membranes to proximity and drive the merger of only the proximal monolayers. As a result, the membrane and matrix layer deform and locally detach. The fusion site is axially-symmetric; its cross-section is illustrated in Figure 4B. The two fusing membranes form a junction in the center of the stalk, with the two membrane mid-planes forming a corner with a 45° angle. As a result, the lipid tails are sheared and splayed to prevent voids in the hydrocarbon tail moiety^93^. The shear and splay magnitude decays within several nanometers from the stalk and smoothly connects to the flat surrounding membranes. After the stalk has formed, it radially expands to an equilibrium radius *R_D_* by bringing the two inner monolayers into contact along a joint mid-plane, a state called hemifusion diaphragm^94,95^. The rim of the diaphragm is the three-way junction between the diaphragm and the two fusing membranes. The lipid monolayer deformations are continuous; therefore, we explicitly require that the magnitude of lipid splay, saddle-splay, and shear are continuous everywhere in our numerical calculations. The matrix layer adheres to the membrane by electrostatic interaction, and it can locally detach from it at the vicinity of the stalk and the diaphragm rim junction to avoid substantial deformation there. Thus, the matrix is not necessarily parallel to the membrane and can bend independently. The deformation of both the membranes and the matrix layer vanishes at the edge of the fusion site and connects smoothly to a flat surrounding membrane and matrix layer. The membrane fluidity in the lateral direction allows the matrix layer to slide on it freely as the fusion process progresses. The fusion reaction ends by opening and expending a membrane pore within the diaphragm, which must involve the detachment of the favorable bounds between the EBOV luminal monolayer and the VP40 matrix layer.

#### The lipid membrane

We model the lipid membrane using the well-established theory of lipid tilt, splay, and saddle-splay^43,44^. The membrane is composed of two monolayers that share a joint mid-plane. The orientation of the lipids in the two monolayers is independent and is given by the lipid director vector, 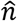. The lipid tilt vector, 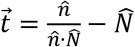, characterizes the shear magnitude and its direction^96^, with 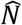 the midplane normal. The monolayer dividing plane is parallel to the membrane midplane and is located at a distance of 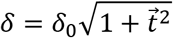 from it, with *δ*_0_ the length of the undeformed monolayer tails. The lipid splay and saddle splay are derived from the splay tensor, 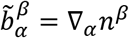, where the sub- and superscripts denote, respectively, the co- and contravariant components in the local coordinate basis of the monolayer dividing plane^44^. Lipid splay is the trace of the splay tensor 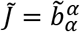, and lipid saddle-splay is its determinant it 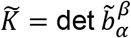. The energy density with respect to the flat tilt-free configuration associated with these deformations is given by^44,97^,

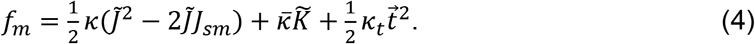

The bending rigidity of the monolayer, *κ_m_*, has a typical value of 10 k_B_T^98^, the saddle-splay modulus, 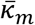, and tilt modulus, *κ_t_* cannot be directly measured and are indirectly estimated. The ratio between saddle-splay modulus and bending rigidity is between −1 to 0^97,99^. The ratio between the bending rigidity to tilt modulus gives a typical tilt decay length of 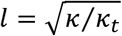, typically between 1 to 2 nm^100^. Here we use *l* = 1.5 nm and 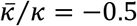. The monolayer spontaneous curvature, *J_sm_*, is the averaged intrinsic curvature of its constituting lipids,

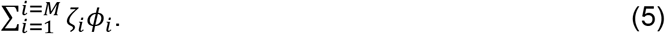

With *M* the total number of lipid components, *ζ_i_*, and *ϕ_i_* are the intrinsic curvature and mole fraction of the *i* lipid components, respectively. The lipid composition is found using lipidomic data of the endosomal and viral membranes (Fig. 1 J). The intrinsic curvature of the most abundant lipids are *ζ_PC_* ≈ −0.1 nm^−1^ for phosphatidylcholine (PC)^101,102^, cholesterol *ζ_CHOL_* ≈ −0.5 nm^−1 101,103^, phosphatidylethanolamine (PE) with *ζ_PE_* ≈ −0.35 nm^−1 103,104^ and sphingomyelin *ζ_SM_* ≈ −0.1 nm^−1 104^. We find that the endosomal and Ebola virus both have monolayer spontaneous curvature of roughly *J_sm_* = −0.22 nm^−1^.

The overall membrane deformation energy is given by the integration of Eq. 4 over the area of both monolayers independently,

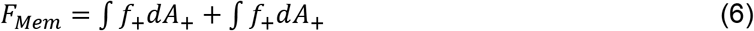

The first and second integrals in Eq. 6 are performed over the upper and lower monolayers area, respectively.

#### The matrix layer

We model the matrix layer as a thin, uniform rigid elastic shell with a flat resting configuration. The matrix can avoid the sharp corners in the vicinity of the stalk and diaphragm rim by local detachment from the membrane. These allow the matrix to avoid strong shear deformations. The elastic energy of matrix deformation up to quadratic order in the area strain, *ϵ*, and in principle curvatures, c_1_ and c_2_, is given by^105^,

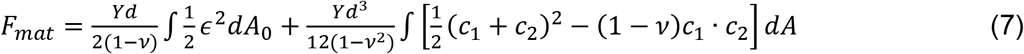

With *dA*_0_ and *dA* the area elements of the undeformed and deformed states, *d* the matrix thickness, *Y* Young’s modulus, *ν* the Poisson’s ratio. We consider only stretching and bending deformations and explicitly prohibit shear. The thickness of the VP40 matrix layer is estimated to be *d* = 4 nm based on the cryo-EM tomography (Fig. 1 A-H). The Young’s modulus and Poisson ratio of the VP40 matrix layer was never measured, but we estimate them to be within the same magnitude as other viruses with similar matrix layer structures, such as M1 of Influenza virus. The VP40 matrix layer Poisson’s ratio is taken as *ν* = 0.5, and the Youngs modulus is in the range 5-22 MPa^106,107^. With that, we estimate the stretching modulus of the VP40 matrix layer in the range of 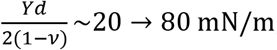, and the pure bending contribution with modulus in the range of 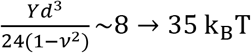.

The matrix layer and the membrane can locally detach in the vicinity of the stalk and diaphragm rim to avoid substantial deformation there. Besides these regions, the matrix interacts continuously with the membrane since the VP40 matrix layer is tightly packed. Inspired by the MD simulations (Fig. 2E), we describe the VP40-membrane interaction energy density with Lennard-Jones-like potential,

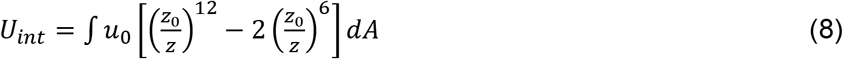

With the integral performed on the area of the matrix layer, *dA*. *z* is the distance from the monolayer dividing plane to the mid-plane of the VP40 layer, *z*_0_ = 4 nm is the resting length obtained from sub-tomogram averaging and the MD simulations (Fig. 4 H). The interaction energy density, *u*_0_, is estimated from the MD simulations as the free energy of a single VP40 dimer at *z* = *z*_0_ (11 k_B_T for pH 7.4 and 6.5 k_B_T for pH 4.5, Fig. 2 E) divided by the density of VP40 dimers obtained from the cryo-EM data (Figure 1 I), we find *u*_0_ = 0.2 k_b_T · nm^−2^ at pH 7.4 and *u*_0_ = 0.1 k_b_T · nm^−2^ at pH 4.5.

#### Way of computation

Our computational approach is based on many previous works^95,108^ and published as open-source code on GitHub (https://github.com/GonenGolani/Fusion_Solver), where further details can be found. The calculation involves three parts – we start by simulating the stalk shape and find its minimal energy configuration. Next, we allow the stalk to expand to the hemifusion diaphragm, and finally, we calculate the energy barrier of pore formation based on the membrane stress and the interaction energy with the VP40 matrix layer in the diaphragm.

The stalk energy barrier represents the minimal mechanical work needed to merge the proximal monolayers. We calculate the hemifusion stalk shape and its formation energy by setting the membrane in stalk configuration. Then, we minimize the sum of the membrane and matrix interaction deformation energies (Eq. 7–8) while requiring that *R_D_* = 0,

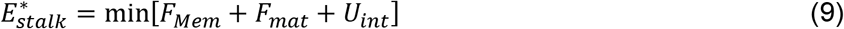

After the stalk has formed, we release the constrain on *R_D_* and allow the system to spontaneously relaxes to a hemifusion diaphragm. The matrix layer can remain attached to the diaphragm or detached.

We calculate the fusion-pore formation energy barrier based on the stress in the hemifusion diaphragm. To facilitate our computation, we assume that the pore formation is initiated at the center of the diaphragm and that the fast fluctuation in pore size does not change the hemifusion diaphragm and matrix layer equilibrium shapes. The pore must expand to the majority of the diaphragm before it overcomes the critical energy, so the initiation point is mainly irrelevant to the magnitude energy barrier. However, since the pores are more likely to form in the vicinity of the diaphragm rim, where stress is maximal, our estimation should be considered a slight overestimation of the actual energy barrier. With this assumption, the energy of pore opening to radius *ρ* is thus given by,

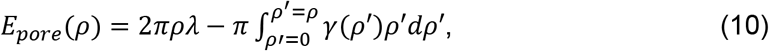

With *λ* the pore rim line-tension magnitude is independent of the matrix layer or the membrane shape. In our simulations, we take it to be *λ* = 12 pN^109^. The second term in Eq. 10 is the energy gained by removing lipids from the stressed diaphragm and relocating them to the surrounding membranes. The stress contains two contributions: the relaxation of the splay, saddle-splay, and shear of the lipids compared to the surrounding membranes and the detachment from the matrix layer,

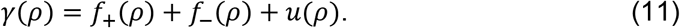

With *f*_+_ and *f*_−_ the energy deformation density of the upper and lower monolayer (Eq. 4), respectively, and *u* the interaction energy density with the matrix (Eq. 8). The pore formation energy barrier is the maxima of *E_pore_* (*ρ*),

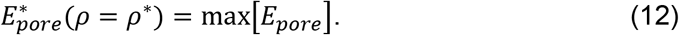

We find the stress (Eq. 11) based on the equilibrium shape of the diaphragm, and Eq. 12 is found by numerically integrating Eq. 10 and finding the maximum.

### Beta-lactamase assay

Huh7 cells were seeded on a 96 well plate coated with 2 μg fibronectin in 1 x PBS at a density of 0.02x 10^6^ cells per well. 24 h post seeding, the media of inhibitor-treated cells was replaced with 25 mM NHCl_4_ in DMEM media (ThermoFisher Scientific) supplemented with 10% (v/v) FBS and 100 U/ml penicillin-streptomycin (ThermoFisher Scientific) and cells were incubated for 1.5 h at 37°C, 5% CO2.

Same amounts of purified beta-lactamase (BlaM)-containing VLPs were either untreated, treated with low pH, thermolysin or a combination of low pH and thermolysin. For the thermolysin-treatment, 500 μg/ml thermolysin (ThermoFisher Scientific), reconstituted in H_2_O and filtered through a 0.22 μm membrane filter, were added to the VLPs for 30 min at 37°C. To quench the reaction, 300 μg/ml phosphoramidon were added for 10 min at 37°C. For the low pH-treatment, citric acid prepared in HNE buffer (10 mM HEPES, 100 mM NaCl, 1 mM EDTA) was added in a final concentration of 1.67 mM to the VLPs for 30 min at 37°C. The BlaM-VLPs were immediately placed on ice until infection.

For infection, the media of all cells was removed, 50 μl pre-treated BlaM-VLP solution were added to each well and the plate was centrifuged for 30 min at 200 g, 20°C (Beckmann). BlaM-VLP solutions were immediately removed and replaced with 100 μl media with and without 25 mM NH_4_Cl. Cells were incubated for 1.5 h at 37°C, 5% CO_2_, before freshly preparing the BlaM dye from the LiveBLAzer™ FRET-B/G Loading Kit with CCF4-AM (ThermoFisher Scientific) supplemented with probenecid (Invitrogen) according to the protocol provided by the manufacturer. 20 μl of the BlaM mix were added per well. After incubation at 11°C for 12-14 h, the cells were briefly checked for viability using a Nikon microscope and detached for 5-10 min using trypsin-EDTA at 37°C. Cells were harvested and washed with 3 x with PBS before performing FACS using a BD FACS Celesta Cell Analyzer (BD Biosciences).

## Acknowledgements

We thank the Infectious Diseases Imaging Platform (IDIP) at the Center for Integrative Infectious Disease Research Heidelberg and the cryo-EM network at the Heidelberg University (HD-cryoNET) for support and assistance; the Electron Microscopy Core Facility at EMBL and Wim Hagen for data acquisition; Dimitrios Papagiannidis for critical reading of the manuscript. Plasmids encoding BlaM-VP40 and pHluorin were a kind gift from Kartik Chandran and Gero Miesenböck, respectively. CSC-IT Centre for Science Ltd. (Espoo, Finland) is acknowledged for excellent computational resources. The authors gratefully acknowledge the data storage service SDS@hd supported by the Ministry of Science, Research, and the Arts Baden-Württemberg (MWK), the German Research Foundation (DFG) through grant INST 35/1314-1 FUGG and INST 35/1503-1 FUGG.

## Author contributions

**SLW**: Conceptualization; Investigation and formal analysis (cloning of reporter constructs, VLP production, (in situ) cryo-ET of VLPs and EBOV-infected host cells, subtomogram averaging, confocal microscopy, BlaM assay); Visualization; Writing - Original Draft. Writing - Review & Editing. **GG**: Investigation and formal analysis (membrane modelling and theory, membrane permeability theory and analysis); Visualization; Writing - Original Draft. Writing - Review & Editing. **FL**: Investigation and formal analysis (MD simulations design, computational resources, data curation, methodology). **MV**: Investigation (in situ cryo-ET of EBOV-infected host cells); Writing - Review & Editing. **KT**: Investigation (BlaM assay). **SSA**: Investigation (acquisition of FACS data). **CL**: Investigation (lipidomics). **OTF**: Writing – Review & Editing. **BB**: Writing – Review & Editing. **TH**: Investigation (infection of host cells with EBOV, purification of EBOV); Writing – Review & Editing. **WN**: Writing – Review & Editing. **USS**: Supervision; Writing – Review & Editing. **PC**: Conceptualization; Funding acquisition; Supervision; Writing - Original Draft; Writing - Review & Editing.

## Data and materials availability

Electron tomography data were deposited to EMDB (EMD-15268, EMD-15244) and will be available upon publication. Additional data and material related to this publication may be obtained upon request.

## Supplementary Figures

**Fig. S1.**
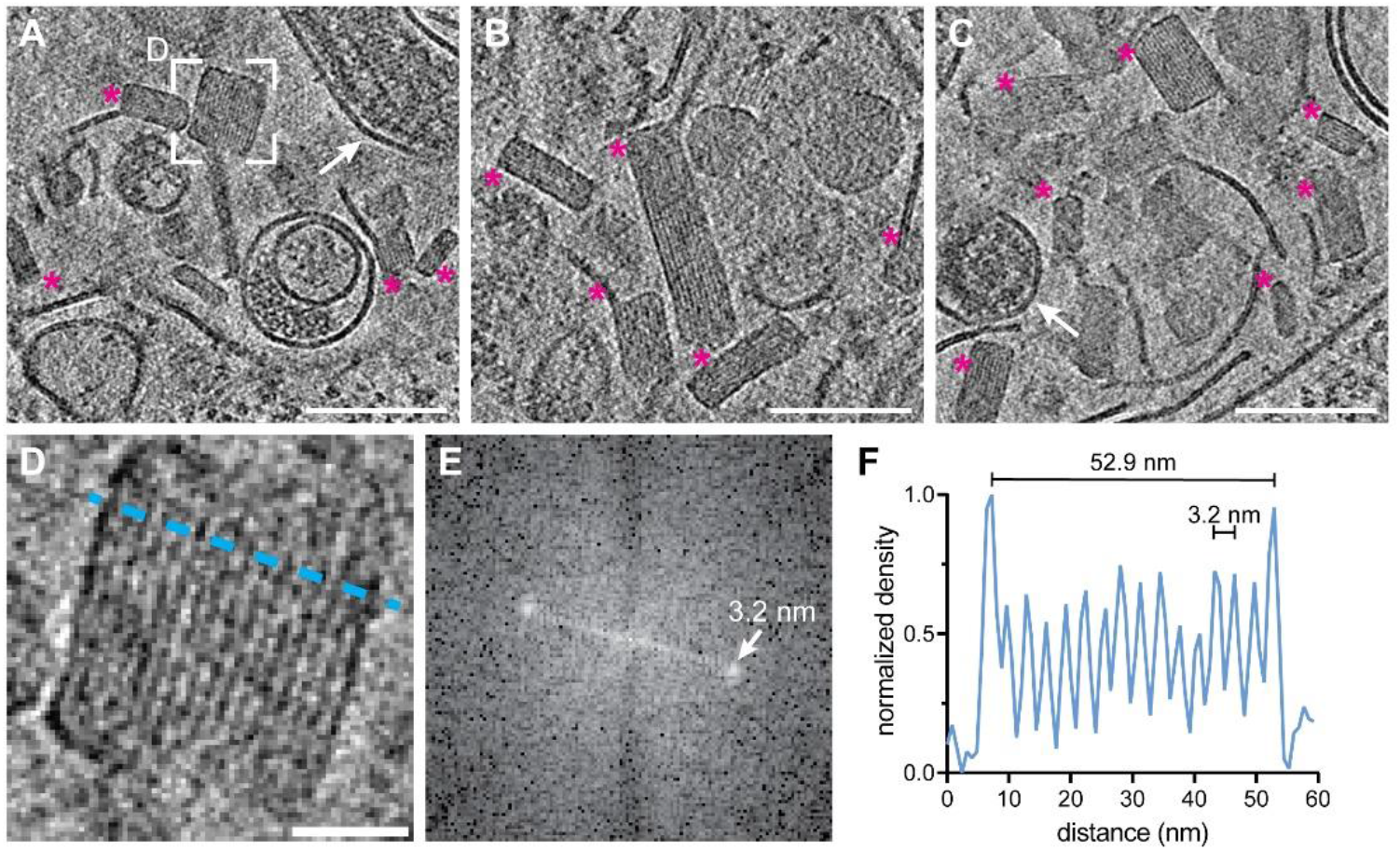
Crystalline lipidic structures in endosomal compartments of EBOV-infected Huh7 cells. **(A-C)** Slices through tomograms showing lumina of endosomal compartments crowded with crystalline lipidic structures (magenta asterisks). Two virions are highlighted with white arrows in (A) and (C). **(D)** Magnified view of the area highlighted in (A) showing a cross-section through a crystalline lipidic structure. To determine the spacing between the stacked lipid monolayers, a line profile was determined (blue line). **(E)** Fourier-transform analysis of the tomogram slice shown in (D) revealing a spacing of 3.2 nm. **(F)** Line profile across the crystal shown in (D) showing the the diameter of the structure along the short axis of 52.9 nm, and the regular 3.2 nm spacing of the lipid monolayers. Scale bars 100 nm (A-C), (D) 20 nm.

**Fig. S2.**
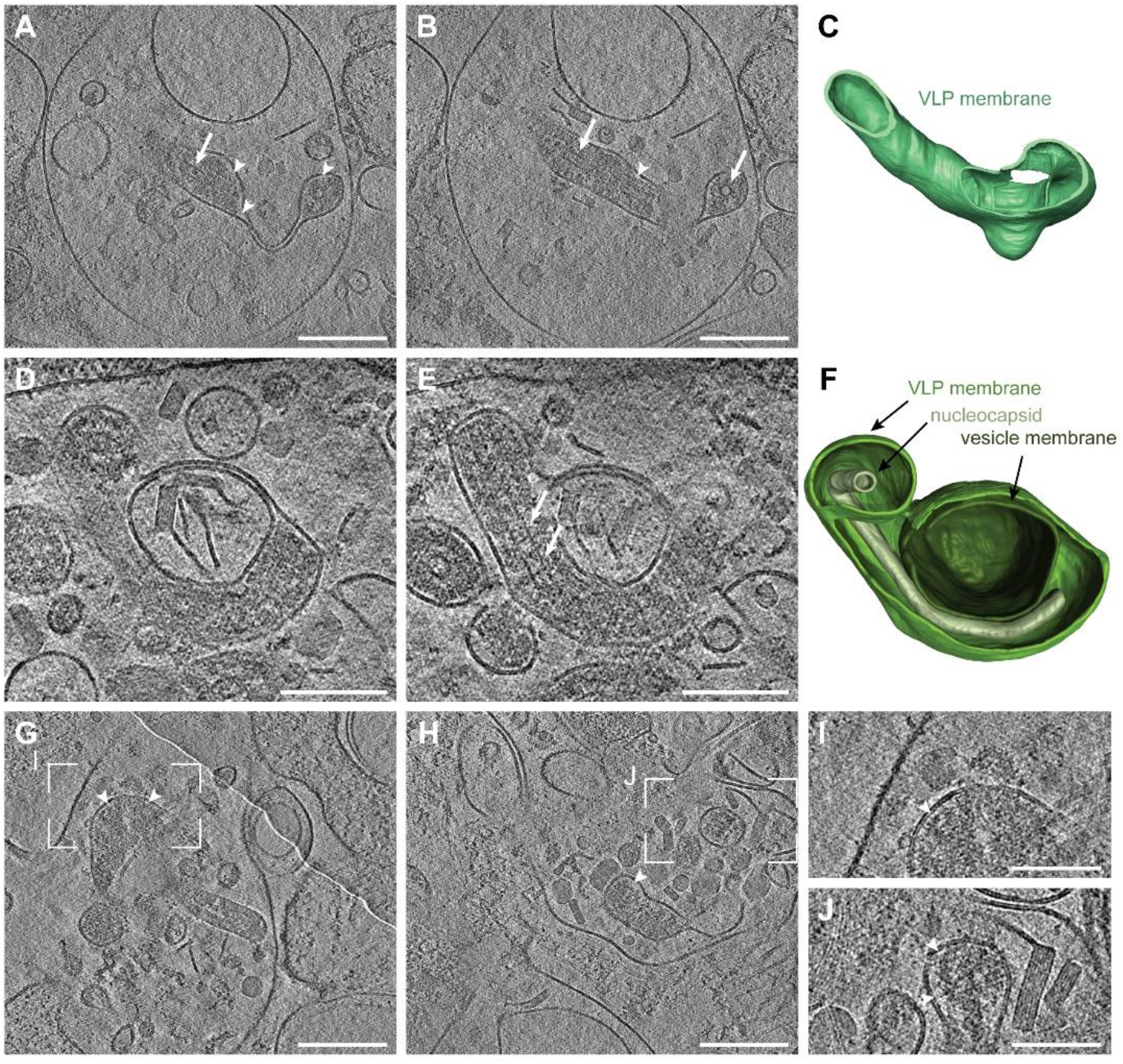
In situ cryo-ET of EBOV infecting Huh7 cells. Slices through tomograms showing Ebola virions inside late endosomal compartments. All virions display condensed nucleocapsids (white arrows) and disassembled VP40 layers, which have detached from the viral membrane as apparent from the gap adjacent to the inner lipid monolayer (white arrowheads). **(A-B)** Different slices through the same tomogram showing an internalized EBOV with a disassembled VP40 matrix and highly flexible membrane. The nucleocapsid is still condensed (white arrow). **(C)** 3D segmentation of the malleable lipid envelope of the EBOV shown in (A) and (B). **(D-E)** Different slices through the same tomogram showing an internalized EBOV with a disassembled VP40 matrix and condensed nucleocapsid. The virion had engulfed an intraluminal vesicle containing cholesterol ester crystals, indicating that this virus has undergone fusion. **(F)** 3D segmentation of EBOV shown in (D) and (E) showing the viral membrane (green), nucleocapsid (light green) and vesicle membrane (dark green). Scale bars: (A), (B) 200 nm, (D-J): 100 nm.

**Fig. S3.**
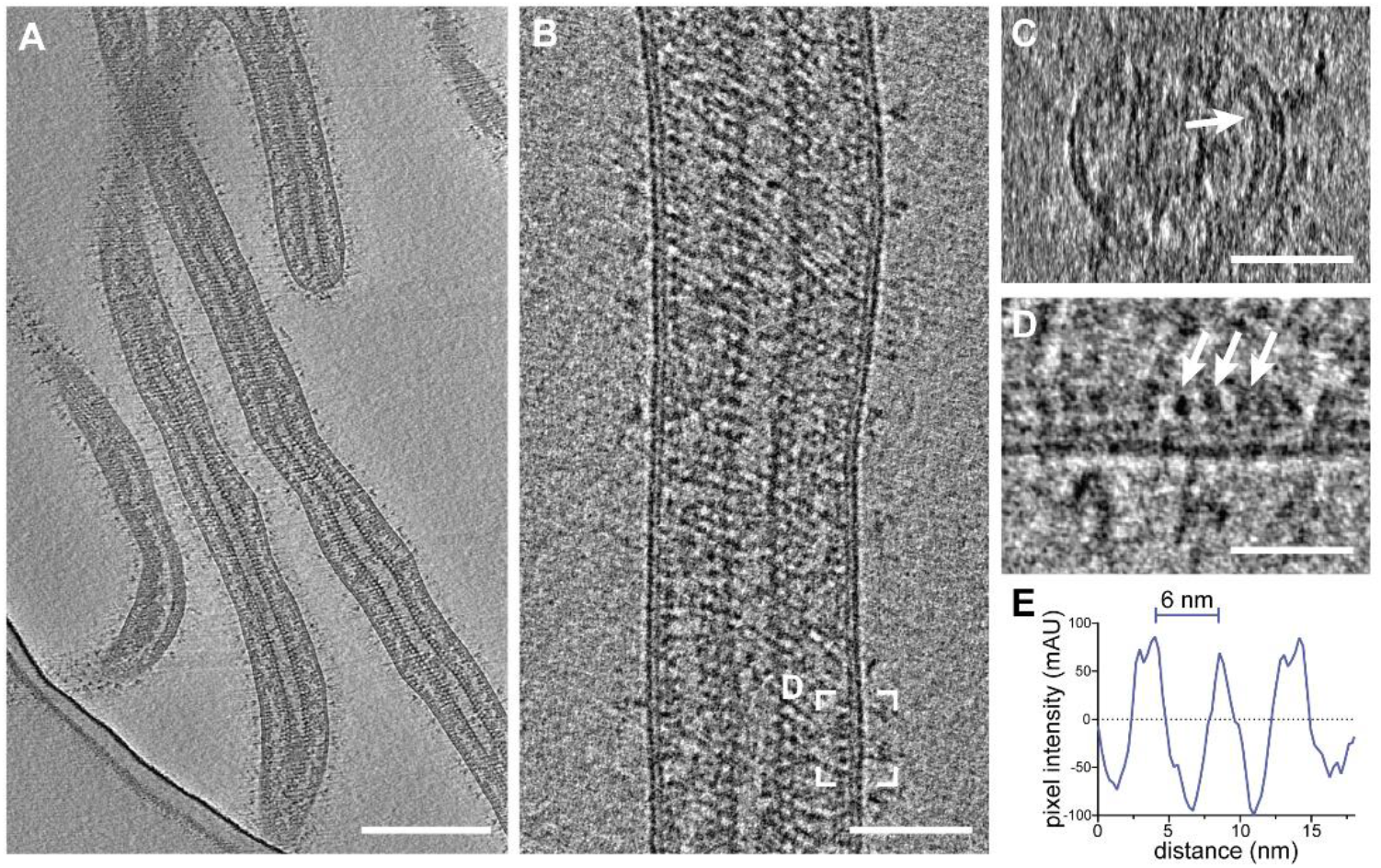
Cryo-electron tomography of purified and chemically fixed EBOV. **(A)** Slices through a tomogram showing an overview of filamentous virions. Condensed and decorated nucleocapsids span the length of each virion. **(B-C)** Longitudinal and transverse cross-section, respectively, of a tomogram containing a filamentous EBOV. **(C)** Transverse cross-section of the virion shown in (B). The VP40 matrix adjacent to the inner membrane monolayer is highlighted by a white arrow. **(D)** Area highlighted in (B) showing a longitudinal cross-section at higher magnification to highlight the VP40 densities lining the inner membrane monolayer. **(E)** Line profile determined adjacent to the inner monolayer of the virion shown in (B) showing the approximately 6 nm pitch of the VP40 matrix. Scale bars: (A), (B): 200 nm, (C): 50 nm, (D): 20 nm.

**Fig. S4.**
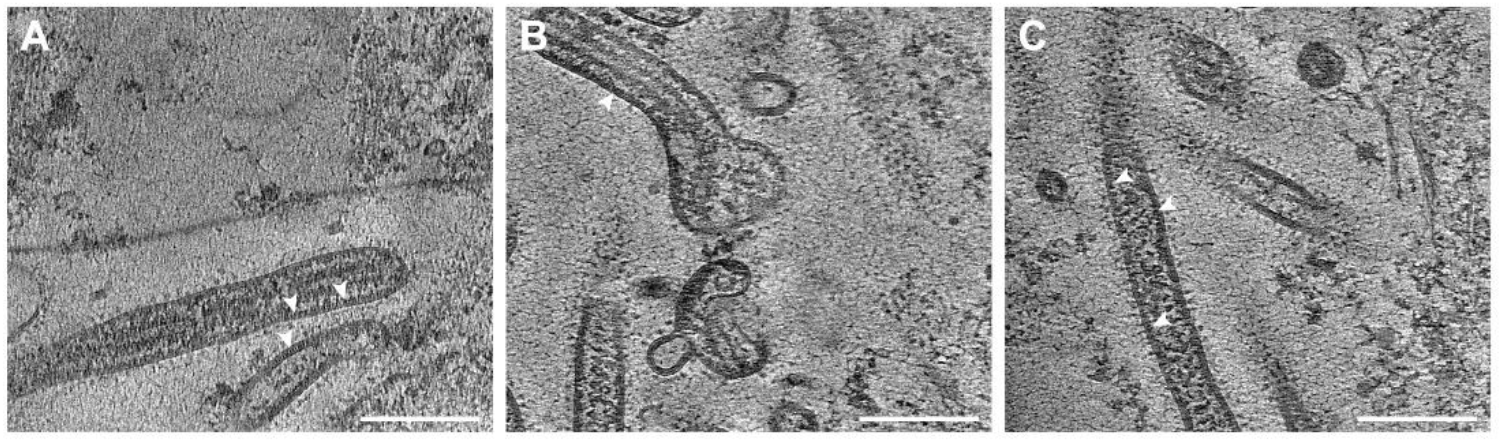
In situ cryo-ET of budding and released EBOV from infecting Huh7 cells. **(A-C)** Slices through tomograms showing Ebola virions adjacent to the plasma membrane of infected Huh7 cells. All virions contain assembled VP40 layers as apparent from the regular densities decorating the inner lipid monolayer at the luminal side (white arrowheads). Scale bars: 200 nm.

**Fig. S5.**
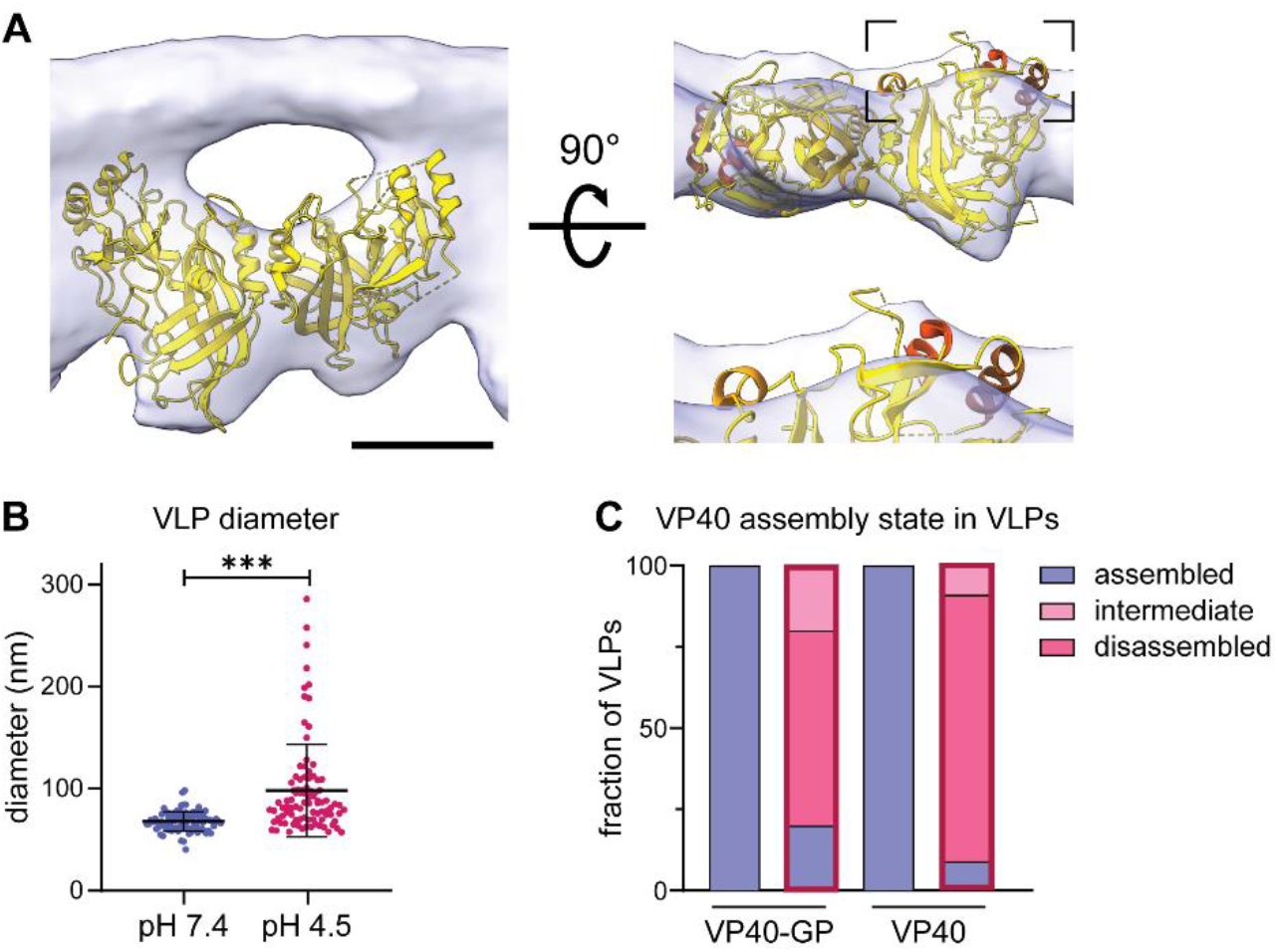
Structural characterization of the VP40 matrix in VLPs by cryo-ET. **(A)** VP40 dimer structure (pdb: 7jzj) fitted into the subtomogram average presented in Fig. 2 from the side view including the density of the inner VLP monolayer and rotated by 90°. Helical segments protruding from the subtomogram average are highlighted in shades of orange. **(B)** Diameter of VLPs composed of GP and VP40 measured from membrane-to-membrane after incubation at pH 7.4 and pH 4.5. Asterisks indicate statistical significance as judged by a two-tailed Welch’s t-test, assuming unequal variance (p<0.0001). **(C)** Quantification of the VP40 assembly state of VLPs composed of VP40 and GP (n= 37 at pH 7.4 and 18 at pH 4.5); or VP40 alone (n= 22 at pH 7.4 and 8 at pH 4.5). The VP40 matrix was either assembled (blue), attached to parts of the VLP membrane (intermediate, pink) or disassembled (dark pink). VP40 assembly in VLPs subjected to neutral or low pH (bars with red frame) was assessed by cryo-ET. Scale bars: (A) 2.5 nm.

**Fig. S6.**
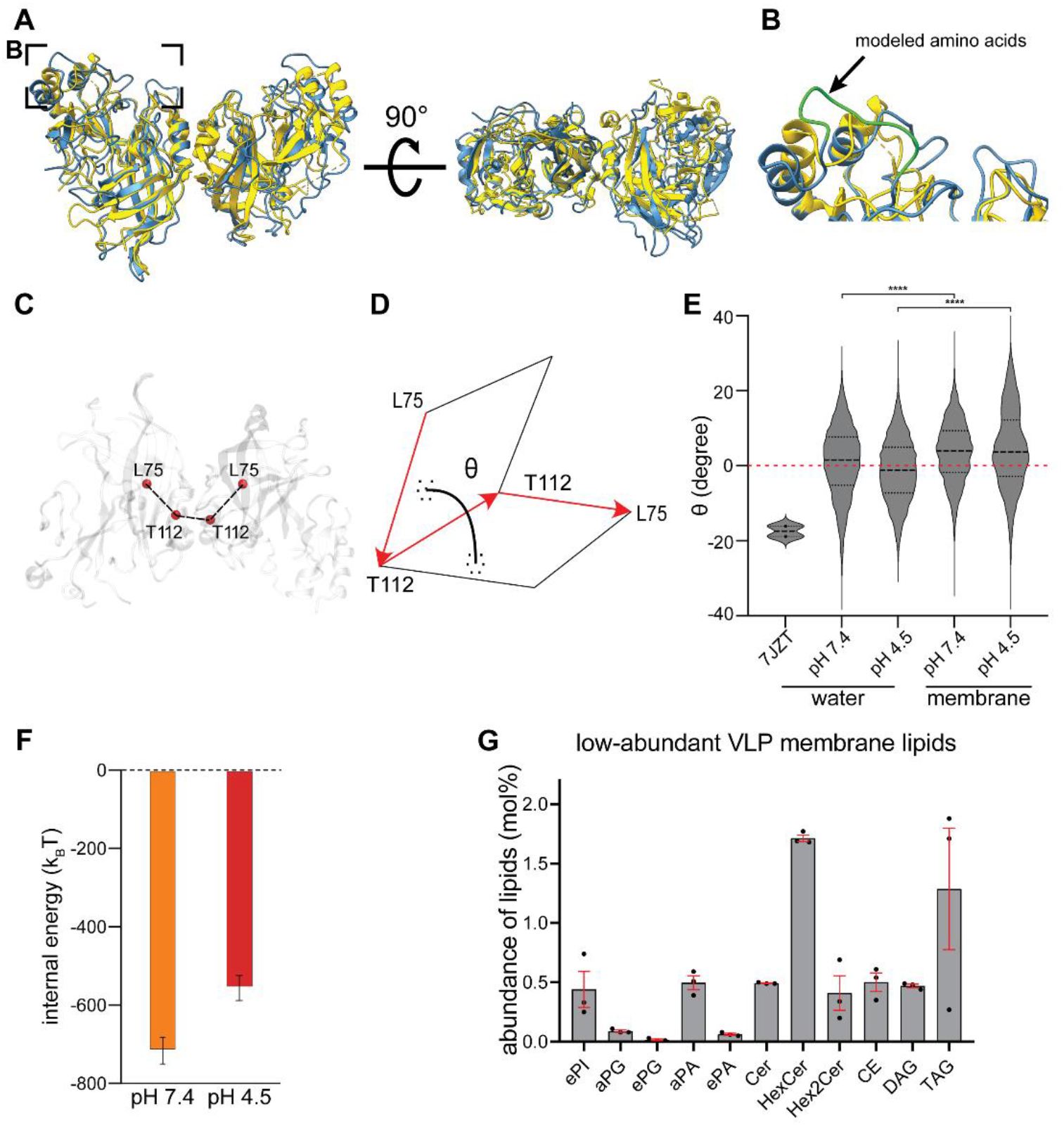
Characterization of VP40 dimer angles and lipidomics. **(A)** Superimposition of the crystallographic structure (pdb: 7jzj, yellow) with the membrane bound VP40 structure from the MD simulations (blue). **(B)** The area highlighted in (A) shows the missing CDTs residues computationally modeled (green). **(C)** Representation of the rotation angle of VP40 monomers along the NTD-dimerization domain. **(D)** The dihedral angle between VP40 monomers is defined as the angle between the plane containing the vector connecting alpha carbon atoms of L75monomer1 and T112monomer1 and the vector connecting atoms T112monomer1 and T112monomer2 and the plane containing this second vector and the vector connecting atoms T112monomer2 and L75monomer2. **(E)** Dihedral angle distribution shows that, regardless of pH, VP40 monomers within the dimer are flexible with a rotation angle oscillating around 1° (SD 9.5) in water, which is 17° smaller than the one measured for the crystallographic structure (pdb: 7jzj). VP40 dimer flexibility is not constrained upon binding to the membrane. However, after binding to the bilayer, the angle distribution was significantly (p ≤ 0.0001) shifted to a value of 3.7° and 4.5° at pH 7.4 and 4.5, respectively. Unpaired t-tests were performed to evaluate the significance of differences in angle distributions. **(F)** VP40-membrane internal energy calculated as the sum of short-range Coulomb and Lennard-Jonson interactions at neutral (orange) and low (red) pH. The internal energies have been calculated over the last 100 ns of the biased MD simulation windows centered at the membrane distance where the free energy minima were reconstructed (3.0 nm and 2.7 nm for neutral and low pHs, respectively). The error was estimated using the block averages over 5 blocks. **(G)** Abundance of low-abundant lipids in the envelope of Ebola VLPs composed of GP, VP40, NP, VP24 and VP35. The mean abundance in mol% and values for each experiment (n=3) are plotted together with the standard error of the mean (red). phosphatidylinositol (PI), phosphatidylglycerol (PG), phosphatidic acid (PA), ceramide (Cer), hexosylceramide (HexCer), cholesterol ester (CE) diacylglycerol (DAG), triacylglycerol (TAG). Prefix “a” indicates acyl-linked glycerophospholipids, prefix “e” indicates ether-linked (plasmanyl) or the presence of one odd and one even chain fatty acyl.

**Fig. S7.**
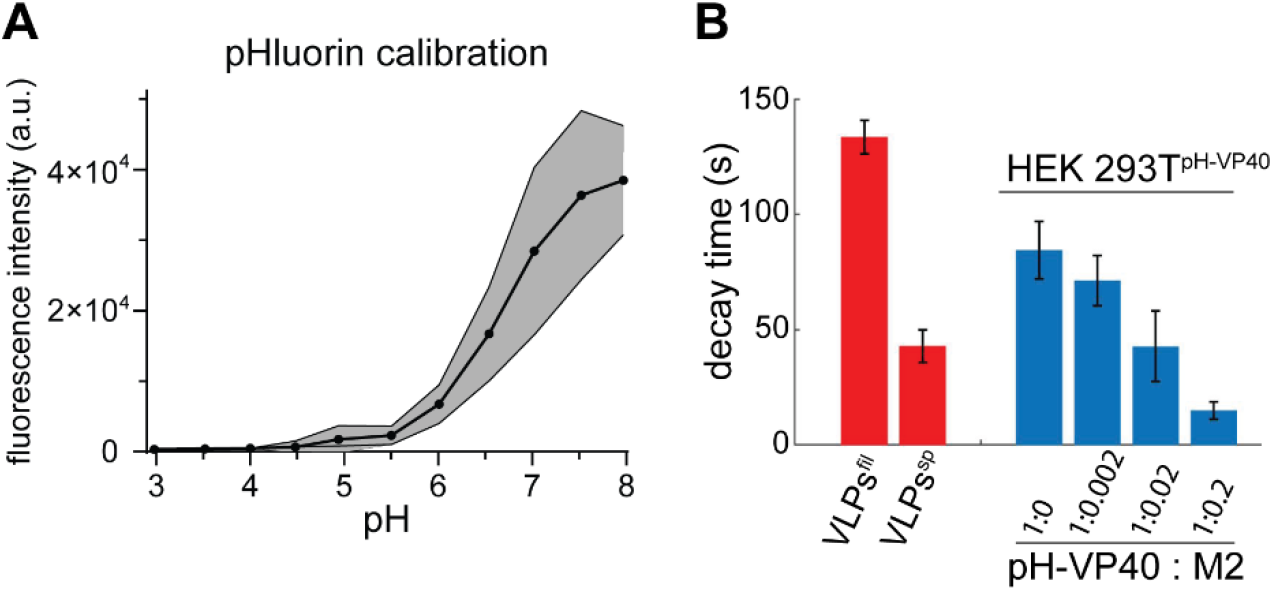
Calibration of pHluorin fluorescence and decay times. **(A)** Fluorescence intensity of HEK 293T cells expressing pHluorin-VP40 measured at 488 nm as a function of pH. Cells were grown in cell culture media before exchanging the media with HNE buffer (10 mM HEPES, 100 mM NaCl, 1 mM EDTA) at different pH. **(B)** pH characteristic decay times as found by fitting the pH levels to Eq. 3 of VLPs (red) and HEK 293T cells (blue) expressing pHluorin-VP40 and influenza M2 in increasing M2 levels. Filamentous VLPs 134±7 sec (n=154), spherical VLPs 43±7 sec (n=66), cells expressing VP40 only 84±12 sec (n=44), cells expressing VP40 and M2 at 1:0.002 molar ratio 71±11 sec (n=30), cells expressing VP40 and M2 at 1:0.02 molar ratio 43±15 sec (n=28), cells expressing VP40 and M2 at 1:0.2 molar ratio 15±4 sec (n=26).

**Fig. S8.**
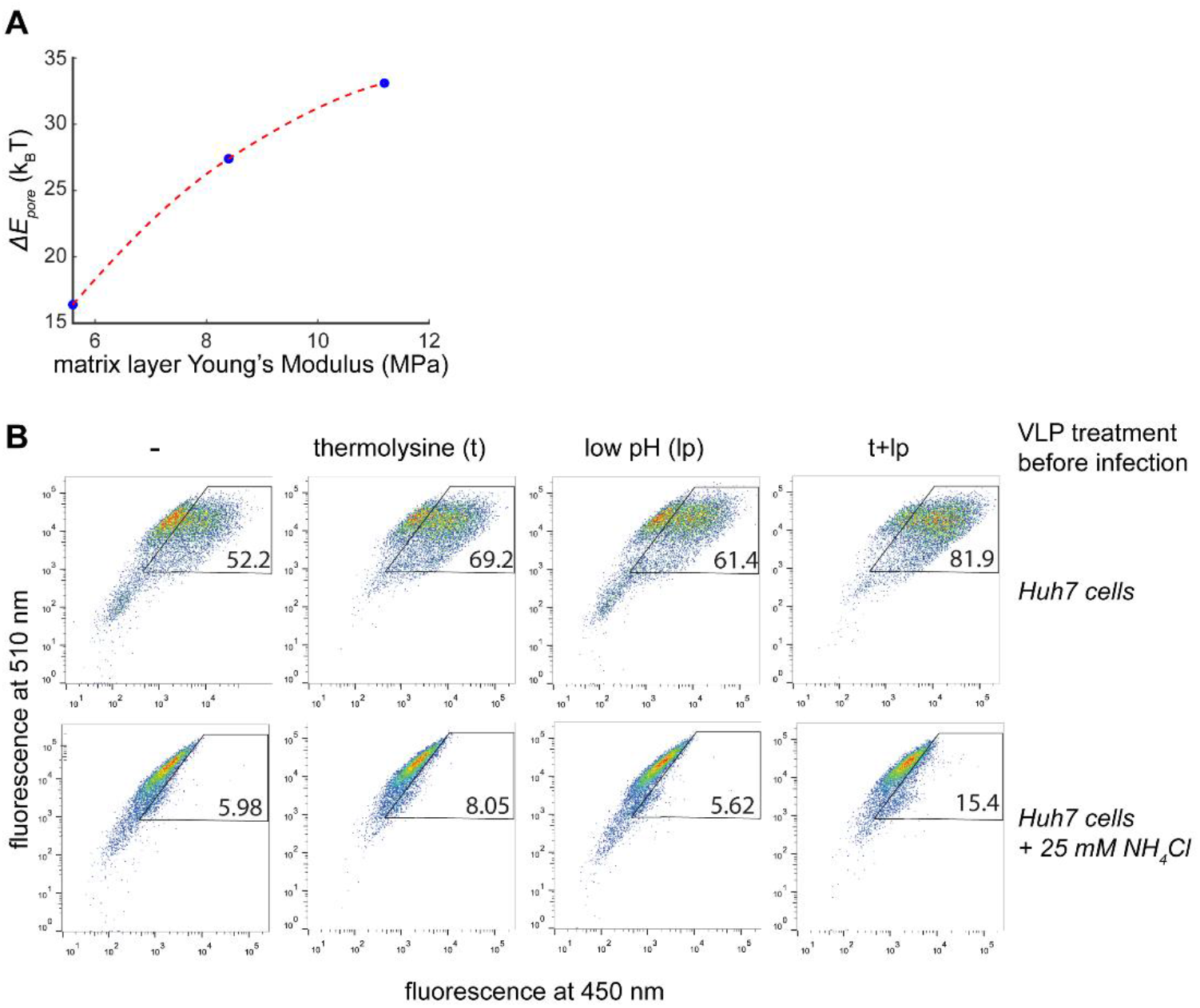
Entry of Ebola BlaM-VLPs into Huh7 cells. **(A)** Plot showing the dependence of the change in fusion pore formation energy between u_0=38 k_B T/nm^2 to u_0=0 (ΔE_pore) on the matrix layer Young’s modulus. Dotted lines serve as a guide to the eye. **(B)**FACS plots showing virus entry as measured by a fluorescence shift of infected cells from emission at 510 nm (no entry) to 450 nm (entry). The top row indicates in vitro VLP treatments prior to infection including buffer control (−), thermolysine treatment, low pH treatment, and a combination of thermolysine and low pH. The first row of FACS data shows entry into Huh7 target cells, the second row shows entry into Huh7 cells treated with 25 mM ammonium chloride to neutralize acidic compartments and assess virus entry in the absence of acidification. FACS data are shown from one out of three repetitions, with 10000 cells measured per sample.

## References

1. Feldmann, H. & Klenk, H. D. Marburg and Ebola viruses. Advances in virus research 47, 1–52 (1996).

2. Geisbert, T. W. & Jahrling, P. B. Differentiation of filoviruses by electron microscopy. Virus Res. 39, 129–150 (1995).

3. Bharat, T. A. M. et al. Structural dissection of Ebola virus and its assembly determinants using cryo-electron tomography. Proc. Natl. Acad. Sci. 109, 4275–4280 (2012).

4. Amiar, S. et al. Lipid-specific oligomerization of the Marburg virus matrix protein VP40 is regulated by two distinct interfaces for virion assembly. J. Biol. Chem. 296, (2021).

5. Johnson, K. A., Taghon, G. J. F., Scott, J. L. & Stahelin, R. V. The Ebola Virus matrix protein, VP40, requires phosphatidylinositol 4,5-bisphosphate (PI(4,5)P 2) for extensive oligomerization at the plasma membrane and viral egress. Sci. Rep. 6, 1–14 (2016).

6. Gc, J. B., Gerstman, B. S., Stahelin, R. V. & Chapagain, P. P. The Ebola virus protein VP40 hexamer enhances the clustering of PI(4,5)P2 lipids in the plasma membrane. Phys. Chem. Chem. Phys. 18, 28409–28417 (2016).

7. Noda, T. et al. Ebola virus VP40 drives the formation of virus-like filamentous particles along with GP. J. Virol. 76, 4855–65 (2002).

8. Wan, W. et al. Ebola and marburg virus matrix layers are locally ordered assemblies of VP40 dimers. Elife 9, 1–22 (2020).

9. Takamatsu, Y., Kolesnikova, L. & Becker, S. Ebola virus proteins NP, VP35, and VP24 are essential and sufficient to mediate nucleocapsid transport. Proc. Natl. Acad. Sci. U. S. A. 115, 1075–1080 (2018).

10. Wan, W. et al. Structure and assembly of the Ebola virus nucleocapsid. Nat. Publ. Gr. 551, (2017).

11. Greber, U. F., Singh, I. & Helenius, A. Mechanisms of virus uncoating. (1994).

12. Yamauchi, Y. & Greber, U. F. Principles of Virus Uncoating: Cues and the Snooker Ball. Traffic 17, 569–592 (2016).

13. Lee, J. E. & Saphire, E. O. Ebolavirus glycoprotein structure and mechanism of entry. Futur. Virol. 4, 621–635 (2009).

14. Dube, D. et al. The Primed Ebolavirus Glycoprotein (19-Kilodalton GP 1,2): Sequence and Residues Critical for Host Cell Binding. J. Virol. 83, 2883–2891 (2009).

15. Nanbo, A. et al. Ebolavirus is internalized into host cells via macropinocytosis in a viral glycoprotein-dependent manner. PLoS Pathog. (2010). doi:10.1371/journal.ppat.1001121

16. Brecher, M. et al. Cathepsin Cleavage Potentiates the Ebola Virus Glycoprotein To Undergo a Subsequent Fusion-Relevant Conformational Change. J. Virol. 86, 364–372 (2012).

17. Carette, J. E. et al. Ebola virus entry requires the cholesterol transporter Niemann-Pick C1. Nature 477, 340–343 (2011).

18. Côté, M. et al. Small molecule inhibitors reveal Niemann-Pick C1 is essential for Ebola virus infection. Nature 477, 344–348 (2011).

19. Miller, E. H. et al. Ebola virus entry requires the host-programmed recognition of an intracellular receptor. EMBO J. 31, 1947–1960 (2012).

20. Simmons, J. A. et al. Ebolavirus Glycoprotein Directs Fusion through NPC1-Endolysosomes. J. Virol. 90, 605–610 (2016).

21. Banerjee, I. et al. Influenza A virus uses the aggresome processing machinery for host cell entry. Science (80-.). 346, 473–477 (2014).

22. Li, S. et al. PH-ontrolled two-step uncoating of influenza virus. Biophys. J. 106, 1447–1456 (2014).

23. Lozach, P. Y., Huotari, J. & Helenius, A. Late-penetrating viruses. Curr. Opin. Virol. 1, 35–43 (2011).

24. van Niel, G. et al. Apolipoprotein E Regulates Amyloid Formation within Endosomes of Pigment Cells. Cell Rep. 13, 43–51 (2015).

25. Klein, S. et al. Post-correlation on-lamella cryo-CLEM reveals the membrane architecture of lamellar bodies. (2020). doi:10.1101/2020.02.27.966739

26. Mahamid, J. et al. Liquid-crystalline phase transitions in lipid droplets are related to cellular states and specific organelle association. Proc. Natl. Acad. Sci. U. S. A. 116, 16866–16871 (2019).

27. Wan, W. et al. Structure and assembly of the Ebola virus nucleocapsid. Nat. Int. J. Sci. 551, 394–397 (2017).

28. Wan, W. et al. Ebola and Marburg virus matrix layers are locally ordered assemblies of VP40 dimers. bioRxiv 1–13 (2020).

29. Pastor, R. W. & MacKerell, A. D. Development of the CHARMM force field for lipids. Journal of Physical Chemistry Letters 2, 1526–1532 (2011).

30. Huang, J. & Mackerell, A. D. CHARMM36 all-atom additive protein force field: Validation based on comparison to NMR data. J. Comput. Chem. 34, 2135–2145 (2013).

31. Huang, J. et al. CHARMM36m: An improved force field for folded and intrinsically disordered proteins. Nat. Methods 14, 71–73 (2016).

32. Bornholdt, Z. A. et al. Structural rearrangement of ebola virus vp40 begets multiple functions in the virus life cycle. Cell 154, 763–774 (2013).

33. Tsui, F. C., Ojcius, D. M. & Hubbell, W. L. The intrinsic pKa values for phosphatidylserine and phosphatidylethanolamine in phosphatidylcholine host bilayers. Biophys. J. 49, 459–468 (1986).

34. Stahelin, R. V. Membrane binding and bending in Ebola VP40 assembly and egress. Frontiers in Microbiology 5, (2014).

35. Panchal, R. G. et al. In vivo oligomerization and raft localization of Ebola virus protein VP40 during vesicular budding. Proc. Natl. Acad. Sci. U. S. A. 100, 15936–15941 (2003).

36. Miesenböck, G., De Angelis, D. A. & Rothman, J. E. Visualizing secretion and synaptic transmission with pH-sensitive green fluorescent proteins. (1998).

37. Deamer, D. W. & Bramhall, J. Permeability of lipid bilayers to water and ionic solutes. Chem. Phys. Lipids 40, 167–188 (1986).

38. Deamer, D. W. Proton permeation of lipid bilayers. J. Bioenerg. Biomembr. 19, 457–479 (1987).

39. Chernomordik, L. V., Zimmerberg, J. & Kozlov, M. M. Membranes of the world unite! J. Cell Biol. 175, 201–207 (2006).

40. Harrison, S. C. Viral membrane fusion. Nature Structural and Molecular Biology 15, 690–698 (2008).

41. Jahn, R. & Grubmüller, H. Membrane fusion. Curr. Opin. Cell Biol. 14, 488–495 (2002).

42. Chernomordik, L. V. & Kozlov, M. M. Mechanics of membrane fusion. Nat. Struct. Mol. Biol. 2008 157 15, 675–683 (2008).

43. Helfrich, W. Elastic Properties of Lipid Bilayers: Theory and Possible Experiments. Zeitschrift fur Naturforsch. - Sect. C J. Biosci. 28, 693–703 (1973).

44. Hamm, M. & Kozlov, M. M. Elastic energy of tilt and bending of fluid membranes. Eur. Phys. J. E 3, 323–335 (2000).

45. Jones, D. M. & Padilla-Parra, S. The β-lactamase assay: Harnessing a FRET biosensor to analyse viral fusion mechanisms. Sensors (Switzerland) 16, (2016).

46. Stauffer, C. E. The effect of pH on thermolysin activity. Arch. Biochem. Biophys. 147, 568–570 (1971).

47. Mingo, R. M. et al. EBOV and SARS-CoV Display Late Cell Entry Kinetics: Evidence that Transport to NPC1 + Endolysosomes Is a Rate-Defining Step. J. Virol. 89, 2931–2943 (2015).

48. Del Vecchio, K. et al. A cationic, C-terminal patch and structural rearrangements in Ebola virus matrix VP40 protein control its interactions with phosphatidylserine. (2018). doi:10.1074/jbc.M117.816280

49. Bornholdt, Z. A. et al. Structural rearrangement of ebola virus vp40 begets multiple functions in the virus life cycle. Cell 154, 763–774 (2013).

50. Ruigrok, R. W. H. et al. Structural characterization and membrane binding properties of the matrix protein VP40 of Ebola virus. J. Mol. Biol. 300, 103–112 (2000).

51. Lee, J. et al. Ebola virus glycoprotein interacts with cholesterol to enhance membrane fusion and cell entry. Nat. Struct. Mol. Biol. 28, (2021).

52. Soni, S. P. & Stahelin, R. V. The Ebola Virus Matrix Protein VP40 Selectively Induces Vesiculation from Phosphatidylserine-enriched Membranes. J. Biol. Chem. 289, 33590–33597 (2014).

53. Adu-Gyamfi, E. et al. The ebola virus matrix protein penetrates into the plasma membrane: A key step in viral protein 40 (VP40) oligomerization and viral egress. J. Biol. Chem. 288, 5779–5789 (2013).

54. Pavadai, E., Gerstman, B. S. & Chapagain, P. P. A cylindrical assembly model and dynamics of the Ebola virus VP40 structural matrix. Sci. Rep. 8, 9776 (2018).

55. Scianimanico, S. et al. Membrane association induces a conformational change in the Ebola virus matrix protein. EMBO J. 19, 6732–6741 (2000).

56. Scianimanico, S. et al. Membrane association induces a conformational change in the Ebola virus matrix protein. EMBO J. 19, 6732–6741 (2000).

57. Nguyen, T. L. et al. An all-atom model of the pore-like structure of hexameric VP40 from Ebola: Structural insights into the monomer-hexamer transition. J. Struct. Biol. 151, 30–40 (2005).

58. GC, J. B., Gerstman, B. S. & Chapagain, P. P. Membrane association and localization dynamics of the Ebola virus matrix protein VP40. Biochim. Biophys. Acta - Biomembr. 1859, 2012–2020 (2017).

59. Booth, T. F., Rabb, M. J. & Beniac, D. R. How do filovirus filaments bend without breaking? Trends Microbiol. 21, 583–593 (2013).

60. Norris, M. J. et al. Measles and Nipah virus assembly: Specific lipid binding drives matrix polymerization. Sci. Adv. 8, (2022).

61. Fontana, J. & Steven, A. C. At Low pH, Influenza Virus Matrix Protein M1 Undergoes a Conformational Change Prior to Dissociating from the Membrane. J. Virol. 87, 5621–5628 (2013).

62. Manzoor, R., Igarashi, M. & Takada, A. Influenza A Virus M2 Protein: Roles from Ingress to Egress. Int. J. Mol. Sci. 18, (2017).

63. Welsch, S., Kolesnikova, L., Krähling, V., Riches, J. D. & Becker, S. Electron Tomography Reveals the Steps in Filovirus Budding. PLoS Pathog 6, 1000875 (2010).

64. DeCoursey, T. E. Voltage-Gated Proton Channels. Cell. Mol. Life Sci. 65, 2554 (2008).

65. Mingo, R. M. et al. Ebola Virus and Severe Acute Respiratory Syndrome Coronavirus Display Late Cell Entry Kinetics: Evidence that Transport to NPC1 ^+^ Endolysosomes Is a Rate-Defining Step. J. Virol. 89, 2931–2943 (2015).

66. Winter, S. L. & Chlanda, P. Dual-axis Volta phase plate cryo-electron tomography of Ebola virus-like particles reveals actin-VP40 interactions. J. Struct. Biol. 213, 107742 (2021).

67. Klein, S. et al. SARS-CoV-2 structure and replication characterized by in situ cryo-electron tomography. Nat. Commun. 1–10 (2020). doi:10.1101/2020.06.23.167064

68. Hagen, W. J. H., Wan, W. & Briggs, J. A. G. Implementation of a cryo-electron tomography tilt-scheme optimized for high resolution subtomogram averaging. J. Struct. Biol. 197, 191–198 (2017).

69. Mastronarde, D. N. & Held, S. R. Automated tilt series alignment and tomographic reconstruction in IMOD. J. Struct. Biol. 197, 102–113 (2017).

70. Castaño-Díez, D., Kudryashev, M., Arheit, M. & Stahlberg, H. Dynamo: A flexible, user-friendly development tool for subtomogram averaging of cryo-EM data in high-performance computing environments. J. Struct. Biol. 178, 139–151 (2012).

71. Castaño-Díez, D. The Dynamo package for tomography and subtomogram averaging: Components for MATLAB, GPU computing and EC2 Amazon Web Services. in Acta Crystallographica Section D: Structural Biology 73, 478–487 (2017).

72. Coutsias, E. A., Seok, C., Jacobson, M. P. & Dill, K. A. A Kinematic View of Loop Closure. J. Comput. Chem. 25, 510–528 (2004).

73. Jo, S., Kim, T., Iyer, V. G. & Im, W. CHARMM-GUI: A web-based graphical user interface for CHARMM. J. Comput. Chem. 29, 1859–1865 (2008).

74. Søndergaard, C. R., Olsson, M. H. M., Rostkowski, M. & Jensen, J. H. Improved treatment of ligands and coupling effects in empirical calculation and rationalization of p K a values. J. Chem. Theory Comput. 7, 2284–2295 (2011).

75. Jo, S., Lim, J. B., Klauda, J. B. & Im, W. CHARMM-GUI membrane builder for mixed bilayers and its application to yeast membranes. Biophys. J. 97, 50–58 (2009).

76. Parrinello, M. & Rahman, A. Polymorphic transitions in single crystals: A new molecular dynamics method. J. Appl. Phys. 52, 7182–7190 (1981).

77. Hoover, W. G. Canonical dynamics: Equilibrium phase-space distributions. Phys. Rev. A 31, 1695–1697 (1985).

78. Essmann, U. et al. A smooth particle mesh Ewald method. J. Chem. Phys. 103, 8577–8593 (1995).

79. Hess, B., Bekker, H., Berendsen, H. J. C. & Fraaije, J. G. E. M. LINCS: A linear constraint solver for molecular simulations. J. Comput. Chem. 18, 1463–1472 (1997).

80. Abraham, M. J. et al. Gromacs: High performance molecular simulations through multi-level parallelism from laptops to supercomputers. SoftwareX 1–2, 19–25 (2015).

81. Jorgensen, W. L., Chandrasekhar, J., Madura, J. D., Impey, R. W. & Klein, M. L. Comparison of simple potential functions for simulating liquid water. J. Chem. Phys. 79, 926–935 (1983).

82. Humphrey, W., Dalke, A. & Schulten, K. VMD: Visual molecular dynamics. J. Mol. Graph. 14, 33–38 (1996).

83. Torrie, G. M. & Valleau, J. P. Monte Carlo free energy estimates using non-Boltzmann sampling: Application to the sub-critical Lennard-Jones fluid. Chem. Phys. Lett. 28, 578–581 (1974).

84. Torrie, G. M. & Valleau, J. P. Nonphysical sampling distributions in Monte Carlo free-energy estimation: Umbrella sampling. J. Comput. Phys. 23, 187–199 (1977).

85. Hub, J. S., De Groot, B. L. & Van Der Spoel, D. G-whams-a free Weighted Histogram Analysis implementation including robust error and autocorrelation estimates. J. Chem. Theory Comput. 6, 3713–3720 (2010).

86. Bonomi, M. et al. Promoting transparency and reproducibility in enhanced molecular simulations. Nat. Methods 2019 168 16, 670–673 (2019).

87. Tribello, G. A., Bonomi, M., Branduardi, D., Camilloni, C. & Bussi, G. PLUMED 2: New feathers for an old bird. Comput. Phys. Commun. 185, 604–613 (2014).

88. Blight, E. G. & Dyer, W. J. A rapid method of total lipid extraction and purification. Can. J. Biochem. Physiol. 37, 911–917 (1959).

89. Malek, M. et al. Inositol triphosphate-triggered calcium release blocks lipid exchange at endoplasmic reticulum-Golgi contact sites. Nat. Commun. 12, (2021).

90. Özbalci, C., Sachsenheimer, T. & Brügger, B. Quantitative analysis of cellular lipids by nano-electrospray ionization mass spectrometry. Methods Mol. Biol. 1033, 3–20 (2013).

91. Liebisch, G. et al. High throughput quantification of cholesterol and cholesteryl ester by electrospray ionization tandem mass spectrometry (ESI-MS/MS). Biochim. Biophys. Acta - Mol. Cell Biol. Lipids 1761, 121–128 (2006).

92. Fuhrmans, M., Marelli, G., Smirnova, Y. G. & Müller, M. Mechanics of membrane fusion/pore formation. Chem. Phys. Lipids 185, 109–128 (2015).

93. Kozlovsky, Y. & Kozlov, M. M. Stalk model of membrane fusion: Solution of energy crisis. Biophys. J. 82, 882–895 (2002).

94. Kozlovsky, Y., Chernomordik, L. V. & Kozlov, M. M. Lipid intermediates in membrane fusion: Formation, structure, and decay of hemifusion diaphragm. Biophys. J. 83, 2634–2651 (2002).

95. Golani, G. et al. Myomerger promotes fusion pore by elastic coupling between proximal membrane leaflets and hemifusion diaphragm. Nat. Commun. 12, 1–18 (2021).

96. Hamm, M. & Kozlov, M. M. Tilt model of inverted amphiphilic mesophases. Eur. Phys. J. B 6, 519–528 (1998).

97. Terzi, M. M., Ergüder, M. F. & Deserno, M. A consistent quadratic curvature-tilt theory for fluid lipid membranes. J. Chem. Phys. 151, 164108 (2019).

98. Dimova, R. Recent developments in the field of bending rigidity measurements on membranes. Advances in Colloid and Interface Science 208, 225–234 (2014).

99. Templer, R. H., Khoo, B. J. & Seddon, J. M. Gaussian Curvature Modulus of an Amphiphilic Monolayer. Langmuir 14, 7427–7434 (1998).

100. Terzi, M. M. & Deserno, M. Novel tilt-curvature coupling in lipid membranes. J. Chem. Phys. 147, 084702 (2017).

101. Chen, Z. & Rand, R. P. The influence of cholesterol on phospholipid membrane curvature and bending elasticity. Biophys. J. 73, 267–276 (1997).

102. Szule, J. A., Fuller, N. L. & Peter Rand, R. The effects of acyl chain length and saturation of diacylglycerols and phosphatidylcholines on membrane monolayer curvature. Biophys. J. 83, 977–984 (2002).

103. Kollmitzer, B., Heftberger, P., Rappolt, M. & Pabst, G. Monolayer spontaneous curvature of raft-forming membrane lipids. Soft Matter 9, 10877–10884 (2013).

104. Leikin, S., Kozlov, M. M., Fuller, N. L. & Rand, R. P. Measured effects of diacylglycerol on structural and elastic properties of phospholipid membranes. Biophys. J. 71, 2623–2632 (1996).

105. Landau, L. D. & Lifshitz, E. M. Theory of Elasticity. (Pergamon PRess, 1970).

106. Li, S., Eghiaian, F., Sieben, C., Herrmann, A. & Schaap, I. A. T. Bending and Puncturing the Influenza Lipid Envelope. Biophys. J. 100, 637–645 (2011).

107. Schaap, I. A. T., Eghiaia, F., Des George, A. & Veigel, C. Effect of Envelope Proteins on the Mechanical Properties of Influenza Virus. J. Biol. Chem. 287, 41078–41088 (2012).

108. Zucker, B., Golani, G. & Kozlov, M. Model for ring closure in ER tubular network dynamics - BioRxiv. bioRxiv 102–104 (2021). doi:10.4324/9780203619308-15

109. Portet, T. & Dimova, R. A new method for measuring edge tensions and stability of lipid bilayers: Effect of membrane composition. Biophys. J. 99, 3264–3273 (2010).

